# Oregano essential oil vapour prevents *Plasmopora viticola* infection in grapevine (*Vitis Vinifera*) by triggering autoimmune metabolic pathways

**DOI:** 10.1101/602730

**Authors:** Markus Rienth, Julien Crovadore, Sana Ghaffari, François Lefort

## Abstract

The reduction of synthetic fungicides in agriculture a major challenge in maintaining sustainable production, protecting the environment and consumers’ health. Downy mildew caused by the oomycete Plasmopora viticola is the major pathogen in viticulture worldwide and responsible for up to 60% of pesticide treatments. Alternatives to reduce fungicides are thus utterly needed to ensure sustainable vineyard-ecosystems, consumer health and public acceptance. Essential oils (EOs) are amongst the most promising natural plant protection alternatives and have shown their antibacterial, antiviral and antifungal properties on several agricultural crops. However, the efficiency of EOs highly depends on timing, application method and the molecular interactions between the host, the pathogen and EO. Despite proven EO efficiency, the underlying processes are still not understood and remain a black box. The objectives of the present study were: a) to evaluate whether a continuous fumigation of a particular EO can control downy mildew in order to circumvent the drawbacks of direct application, b) to decipher molecular mechanisms that could be triggered in the host and the pathogen by EO application and c) to try to differentiate whether essential oils directly repress the oomycete or act as plant resistance primers.

A custom-made climatic chamber was used for a continuous fumigation of potted vines with different EOs during long-term experiments. The grapevine (Vitis vinifera) cv Chasselas was chosen in reason of its high susceptibility to Plasmopara viticola. Grapevine cuttings were infected with P. viticola. and subsequently exposed to continuous fumigation of different EOs at different concentrations, during 2 application time spans (24 hours and 10 days). Experiments were stopped when infection symptoms were clearly observed on the leaves of the control plants. Plant physiology (photosynthesis and growth rate parameters) were recorded and leaves were sampled at different time points for subsequent RNA extraction and transcriptomics analysis. Strikingly, the Oregano vulgare essential oil vapour treatment during 24h post-infection proved to be sufficient to reduce downy mildew development by 95%. Total RNA was extracted from leaves of 24h and 10d treatments and used for whole transcriptome shotgun sequencing (RNA-seq). Sequenced reads were then mapped onto the V. vinifera and P. viticola genomes. Less than 1% of reads could be mapped onto the P. viticola genome from treated samples, whereas up to 30 % reads from the controls mapped onto the P. viticola genome, thereby confirming the visual observation of P. viticola absence in the treated plants. On average, 80 % of reads could be mapped onto the V. vinifera genome for differential expression analysis, which yielded 4800 modulated genes. Transcriptomic data clearly showed that the treatment triggered the plant’s innate immune system with genes involved in salicylic, jasmonic acid and ethylene synthesis and signaling, activating Pathogenenesis-Related-proteins as well as phytoalexin synthesis.

These results elucidate EO-host-pathogen interactions for the first time and indicate that the antifungal efficiency of EO is mainly due to the triggering of resistance pathways inside the host plants. This is of major importance for the production and research on biopesticides, plant stimulation products and for resistance-breeding strategies

**Author Summary:** The reduction of synthetic plant protection products is a major concern of modern agriculture. The oomycete *Plasmopora viticola* which causes downy mildew in grapevine is amongst the most important grapevine pests and responsible for the dispersion of huge amounts of pesticides in vineyards. Among the evaluated alternatives to reduce or replace synthetic pesticides, plant volatile compounds could represent a sustainable solution. Some plant essential oils (EOs) have already shown antifungal capacities. However, their application is often difficult in terms of the right timing of treatment, degradation, bad rainfastness, mixability and phytotoxicity.

The aim of the present work was to investigate whether the vapour phase, applied by a continuous fumigation of different EOs, might inhibit the development of downy mildew on grapevine, and in case of proven efficiency, to study the induced transcriptomic changes by RNA-sequencing in an attempt to elucidate the underlaying molecular interactions. Our results showed that the vapour phases of *O. vulgare* and *T. vulgaris* were highly efficient against the pathogen. The study of differentially expressed genes indicated that the EO vapour triggered the main mechanisms of the plant’s innate immune system such as PTI (Pattern-Triggered Immunity) and ETI (Effector Triggered immunity).

For the first time these results highlight the effects of EOs vapour on plant genes expression, which is very valuable information for the development of new natural plant protection products, as well as for breeding disease resistant cultivars.

## Introduction

Global food supply is highly dependent on industrial agriculture, which in turn would not be possible without the intensive use of pesticides against fungal diseases and other pests. Responding to consumers’ increasing demands for a sustainable food production implies developing alternatives to conventional synthetic plant protection products. Long-term fungicide applications have consequently led to increased resistances of pathogens and detrimental impacts on ecosystems and humans (1), followed by a decreasing acceptance by consumers (2). This is particularly true for grapevine (*Vitis vinifera* L.), which is highly sensitive to fungal diseases such as downy mildew caused by the obligate biotrophic pathogenic oomycete *Plasmopara viticola* (Berk. & M.A. Curtis) Berl. & De Toni, (1888) which is natural to North America. This organism was accidentally introduced in Europe via infected cuttings at the end of the 19^th^ century and is one of the most devastating diseases of viticulture worldwide (3), which explains that the application of relatively large amounts of pesticides in viticulture, when compared to other crops, is necessary to guarantee yield and quality of grape production.

The infection cycle of *P. viticola* starts with zoomeiospores, which are released by mature zoosporangia germinating from oospores which are the only source of primary inoculum. Encysted zoomeiospores germinate to form a germ tube, which will penetrate the leaf through a stoma. A substomatal vesicle then develops and gives rise to the intercellular mycelium with its many haustoria penetrating cell walls of the mesophyll. The incubation time until visible symptoms appear may last from 4 to approximately 18 days depending on temperature. After this time span, oil-spot lesions appear on the adaxial surface. If the leaf is subsequently incubated in conditions of high humidity, hyphal coils grow into the sub-stomatal cavity and give rise to sporangiophores emerging from the stoma. Incubated in the laboratory at 14-28°C and high humidity (above 90%), the sporangiophores and zoosporangia develop overnight, before liberating zoospores (4).

During the vegetative cycle, growers usually need to treat between 3 to 15 times against downy mildew with systemic or organic fungicides. This represents a fundamental problem regarding the sustainability of ecosystems, biodiversity, consumer health and acceptance as well as long term efficiency of systemic fungicides, along with an increased development of resistances (5, 6)

To reduce synthetic pesticides, organic production is one alternative to conventional farming but still highly dependent on copper (Cu), the oldest and still a very efficient treatment against downy mildew. It remains, however, a heavy metal accumulating in vineyard soils. When compared to the overall average Cu concentration of 16.85 mg kg^−1^ in soils, vineyards have the highest mean soil Cu concentration (49.26 mg kg^−1^) of all land categories, followed by olive groves and orchards (7, 8). Cultivation of disease resistant varieties is certainly one of the most ecological solutions to reduce pesticides and, due to tremendous efforts of international public breeding programmes, a large choice of disease-resistant grape cultivars is nowadays available to growers (9–12). However, the organoleptic quality is still often inferior to the one of traditional cultivars, making them less attractive to consumers and thus producers. For these last reasons, access of organic wines to the wine market is still difficult. Furthermore, the durability of resistance factors is often not stable, in particular in monogenetic cultivars (13). Genetically modified organisms are not a solution either, since they are so far neither premised nor authorised in most producing countries. Alternative plant protection strategies are thus utterly needed to guarantee a sustainable viticulture that is both environment- and consumer-friendly.

It is now widely acknowledged that plants possess two forms of innate immune system responses against pathogens. Molecular expression patterns induced by microbial molecules, i.e. Pathogen, Microbial or Damage-associated molecular pattern (PAMPs, MAMPs or DAMPs) lead to pattern triggered immunity (PTI), which represents the first line of defense of plants against pathogens. PTI leads to a cascade, which is marked by common signaling events, such as ion fluxes, protein phosphorylation cascades, accumulation of reactive oxygen species (ROS), induction of defense genes and cell-wall reinforcement by callose deposition (14, 15).

The second innate immune defense response is the effector-triggered immunity (ETI), where the plant’s response is triggered by pathogen effectors. ETI results from the highly specific, direct or indirect interaction of pathogen effectors and the products of plant disease resistance (R) genes, which leads to a strong local defense response often associated with programmed cell death (PCD) as a part of hypersensitive response (HR) that stops pathogen growth (16).

Inducing resistance to pathogens by activating the plant’s innate immunity through application of natural products which trigger PTI and/or ETI, could thus represent an alternative strategy to protect plants against diseases (17).

Plant phytochemicals have been investigated for decades, and it has been demonstrated that specific plant volatile organic compounds (VOCs) have antifungal, anti-bacterial, as well as repulsive effects on insect pests. It is however not known to what extent VOCs have direct effects on the pathogen or induce stimulation of the host plant’s defense mechanisms.

During the last decade, several different resistance inducers have been tested for their ability to induce defense responses of the susceptible *V. vinifera* against *P. viticola*, such as beta-aminobutyric acid (18) chitosan (19), laminarin, sulfated laminarin (20, 21), Frutogard® and other plant extracts (22, 23).

As natural products, EOs have shown antifungal properties against several pathogens such as *P. viticola, Botrytis cinerea* Pers. 1794 and *Fusarium sp*. (24, 25). The chemical composition and activities of members of the EO-producing family *Lamiaceae* have been widely studied, and the antioxidant activity could mainly be attributed to carnosoic acid, carnosol, rosmarinic acid, and other phenolic compounds, whereas the fungicidal activity seems to be due to molecules such as carvacrol, thymol, and p-cymene (26). The use of EOs to control plant pathogens has been evaluated on different species with different pathogens. In such studies, the timing of EO applications seems to be a crucial point to increase the efficiency of pathogen control. For instance, *Origanum vulgare* L. EO had preventive effects against *B. cinerea* development on tomato leaves, with the highest efficiency when applied 24h post-infection. This suggested an effect on early fungal development such as spore germination, germ tube growth and/or appressorium formation, since light- and electron-scanning microscope observations revealed alterations of hyphal morphology when exposed to EOs (27, 28).

Studies conducted on grapevines, infected with downy mildew and treated with sage (*Salvia officinalis* L.) EO showed a 94% reduction in disease severity. However, due to the very low rainfastness and degradation of EOs, the efficiency in rainy years was much less important (29). This is probably the reason why other field studies on the variety Merlot, using EOs of *Corymbia citriodora* (Hook.)*, Syzygium aromaticum* (L.)*, O. vulgare* and *Thymus vulgaris* L. did not show any efficiency in field trials against downy mildew, whereas thyme and oregano EOs were very efficient in inhibiting fungal growth in Petri dishes. This highlights that a major problem of EO efficiency against pathogens is due to degradation by light, heat, oxygen and humidity (30), application time and bad rainfastness.

Several studies indicate that the vapour phase of EO is more fungitoxic than the contact liquid phase, although this has only been shown for *Botrytis* in Petri dishes (28, 31, 32). Thus a continuous fumigation of plants with EO vapour could possibly circumvent these drawbacks. For a sustainable agricultural production, using EO vapour as a direct treatment with diffusers or in the form of co-plantations of EO-VOCs emitting plants could be considered in integrated systems, able to control fungal diseases.

The mechanisms underpinning the effect of antifungal EO-VOCs are not well understood so far. Some VOCs seem to have a direct effect on pathogens, while others seem to elicit the plant innate immune system with its complex mechanism. Several authors have found genes involved in the biosynthesis of phytoalexins, pathogenesis-related (PR) proteins and cell wall proteins when VOCs were applied. (33). Understanding the mechanism involved in EO efficacy against fungi could thus provide very valuable information when developing natural fungicides, plant defense stimulation products, as well as providing genetic targets for the breeding of resistant varieties.

The aims of the present study were thus to investigate if the vapour phase, applied by a continuous fumigation of different EOs, could inhibit the development of downy mildew on grapevine leaves and, in case of proven efficiency, to study and reveal the induced transcriptomic changes by RNA-sequencing in an attempt to elucidate the underlaying molecular interactions.

## Results & Discussion

### EO vapour impedes *Plasmopora viticola* development

Terpene composition of EOs as determined by GC-FID (Gas Chromatography with Flame-Ionization Detection) analysis, showed that the main constituents were carvacrol (21- 60 %), p- cymene (6- 20 %), γ-terpine (9-26%) and thymol (9-26%) in EO of *O. vulgare* and thymol (24-45%), p-cymene (15-37%) and g-terpinene (6-24%) in *T. vulgaris* (supplemental file S2). These compositions are congruent with the commonly reported values in literature for *O. vulgare* (34–36) and *T. vulgaris* (37). Still, huge variations can occur depending on the species, the collecting season, the geographical position, the collected plant organ and the oil extraction method (38).

Concentrations of oil vapour inside the chamber varied for *O. vulgare* between 0.023 and 0.015 % (heated and non-heated respectively) between 0.1 and 0.06 % for *T. vulgaris*. Concentration of vapour without heating was considered sufficiently high, thus experiments were carried out without the heating system.

In the first series of experiments with *O. vulgare* and *T. vulgaris*, vines were treated continuously for 10d after *P. viticola* infection. The visual assessment after sporulation induction (Fig 1.) showed that both oils were highly efficient in inhibiting *P. viticola* development on leaves and reduced disease severity by up to 98 %. Antifungal efficiency of both oils was comparable and not significantly different. Unfortunately, after a 10d exposure to essential oil vapour (of both oils), vines expressed symptoms of an acute phytotoxicity consisting of browning of young leaves, a decline in growth velocity as well as a reduction in photosynthesis (Fig 2.) and leaf-nitrogen content.

**Figure 1:**
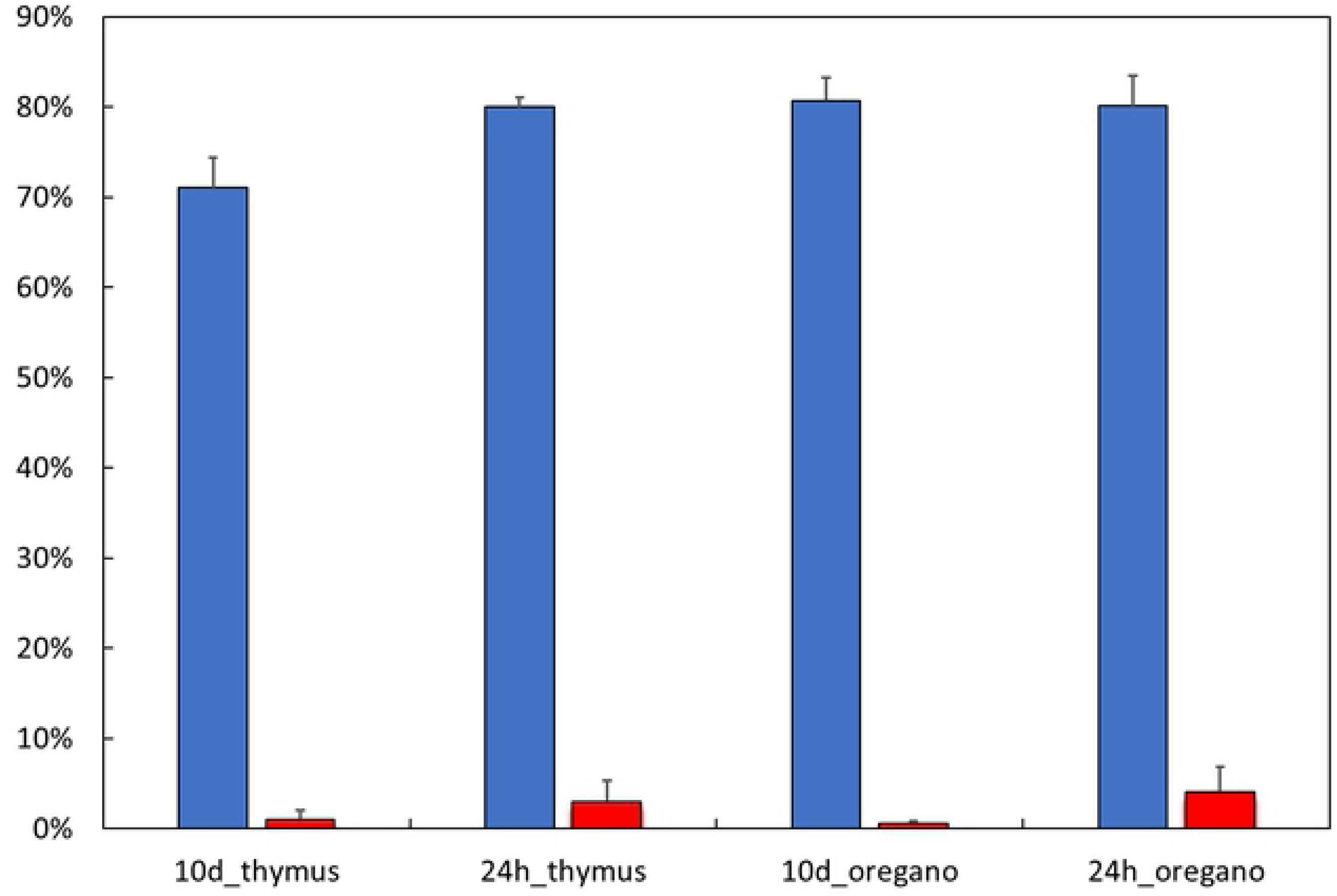
Average Downy mildew severity assessed after sporulation on 6 plants. Blue: non-treated control; red: oregano treatments.

**Figure 2:**
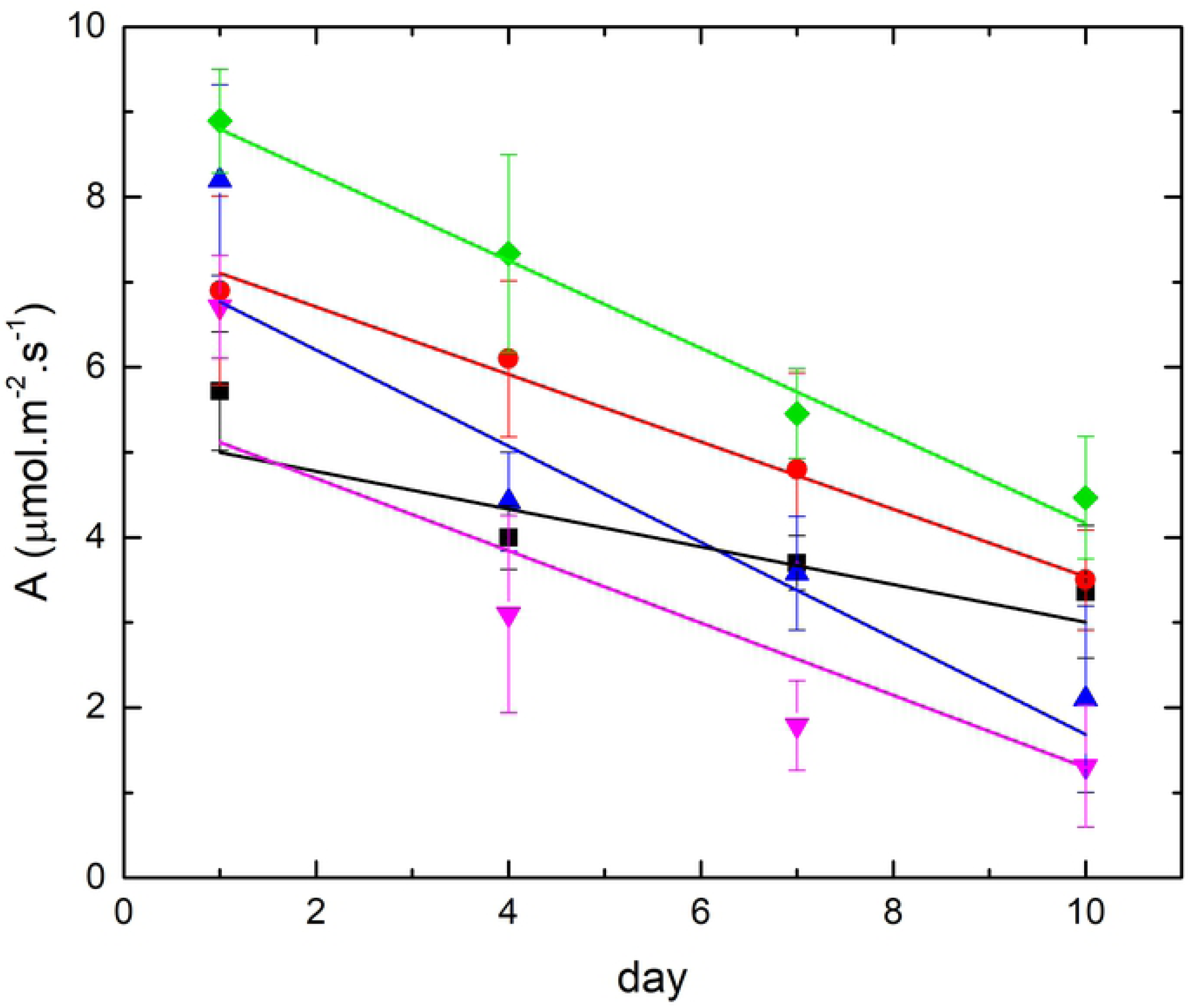
Evolution of photosynthesis of treated and non-treated plant in climatic chamber. Black squares: 24h *T. vulgaris* treatment; red circles 24h *O. vulgare* treatment; blue upward triangles: 10d *O. vulgare* treatment; purple downward triangles: 10d *T. vulgaris* treatment; green diamonds: control.

A second set of experiments was carried out to test an EO treatment time reduced to 24h and still resulting in an antifungal effect, with the advantage of reducing EO quantity and eventually phytotoxicity. For this second test, EO vapour treatments were maintained for only 24h after infection. Infected plants were then kept for 10d in the chamber to assess disease severity after induction of sporulation, as for the first experiments. Interestingly, the efficiency of the EO treatment against downy mildew was only slightly but not significantly lower, compared to the 10d treatment with a 95% decrease in disease severity in treated plants (Fig 1.).

This strongly suggests that the antifungal effect of EOs acts during the early stages of the infection cycle or even prior to infection. However, it remains unclear whether this is because of a direct lethal effect either on the zoospores before producing haustoria or on the haustoria growth before entering the stomata or else due to the inhibition of mycelium growth inside the leaf. An alternative hypothesis is that EO vapour stimulates innate plant immunity by activating PTI and/or ETI, which would impede entrance of the haustoria through the stomata and/or limit mycelium growth inside the leaf.

This ability of different EO against downy mildew has been highlighted for crops other than grapevine. For example in cucumber, castor and clove oils significantly reduced the severity of downy mildew (39). Similarly, sage EO applied at a concentration of around 1% reduced disease severity of downy mildew on cucumber by almost 100% in greenhouses but only 70% in the field (40). This illustrates the problem of rainfastness and degradability of EO in field conditions, thereby justifying the vaporisation approach.

Several studies on grapevine have confirmed that EOs could be an efficient alternative treatment against P. *viticola*. Dagostin et al. (2011) used *Salvia officinalis* EO at concentration of 50 mL/L on potted *Pinot gris* vines in greenhouses, and on *Carbernet Sauvignon* in field trials (29). The efficiency was here also much lower in the field due to previously mentioned reasons.

La Torre et al. (2014) used clove and tea-tree oil on leaf discs and in the field on cv. *Malvasia di Candia* against downy mildew and showed that both EOs controlled the development of downy mildew both in situ and *in vivo* (41). An *in vitro* study of direct and vapour phase application of EOs on *Chardonnay* leaves, the EOs being of cinnamon*, Eucalyptus globulus* Labill., marjoram, tee-tree, peppermint, oregano and thyme, on *P. viticola* showed very good efficiency of all oils, with cinnamon and *Eucalyptus globulus* EOs as the most fungitoxic (32). Applying only the vapour phase, has, up to now, never been tested *in vivo* on *V. vinifera*. Studies on other plants showed that EOs, encapsulated in mesoporous silica and subsequently slowly released as vapour, had direct antifungal properties on *Aspergilus niger* (31). Similar results were obtained by Soylu et al., (2007, 2010), where the vapour phase generally showed a higher efficacy against *Botrytis cinerea* (27, 28).

This direct antimicrobial or antifungal activity of EOs might mainly be caused by the properties of their terpenes/terpenoids, that—due to their highly lipophilic nature and low molecular weight—are capable of disrupting the cell membrane, causing cell death or inhibiting the sporulation and germination of fungi (42). Antifungal properties are generally linked to cell membrane disruption, alteration and inhibition of cell wall formation, dysfunction of the fungal mitochondria, inhibition of efflux pumps and / or ROS production. More specifically, the antifungal activity of carvacrol and thymol have often been attributed to their cell membrane damaging effect, because of their interaction with membrane sterols, in particular with ergosterol. Carvacrol would be able to bind with the sterols of the fungal plasma membrane, which would result in damage to the membrane, conducing to the death of the fungus (43). Thymol seems to affect mycelium morphology, with changes in the localisation of chitin within the hyphae (42, 44).

An exhaustive analysis of scientific literature up to the most recent, does not help to clarify whether EOs trigger the innate immune system of host plants or only have a direct effect on pathogens. To test the hypothesis of EOs being potential primers of plant immunity a transcriptomic analysis was carried out; it is presented in the subsequent chapter.

### Global transcriptomic reprogramming induced by *O. vulgare* vapour treatment

RNA-seq was carried out on the experiments with *O. vulgare*, where the 24h and the 10d samples were sequenced and analysed for differentially expressed genes (DEGs).

A total number of 94 million reads were sequenced from all samples. Read alignment to the *V. vinifera* genome resulted in 70 to 90 % of reads that could be mapped onto the grapevine genome (Fig 3.). A second alignment of reads was performed to the *P. viticola* genome, which has been recently published (45), to check the presence or absence of *P. viticola* genes. For the 10d treatment between 25 to 35 % of reads from control samples mapped to the P. *viticola* genome, while only 1-4 % of reads from the treated plants mapped to the P. *viticola* genome. This strongly demonstrates the absence of *P. viticola* when plants were treated with EO vapour (Fig 3.). Interestingly, no difference in mapped reads was found between control and treatment after 24h, where in both cases only a very low number of reads mapped to the *P. viticola* genome, indicating a very low presence of the oomycete inside the host leaves at this early stage of infection, even in control plants. It can thus not be concluded from these present data whether EO treatment had a direct or indirect inhibitory effect on the *Oomycete* development, since its development even in control plants was too reduced to detect *P. viticola* genes for differential expression analysis.

**Figure 3:**
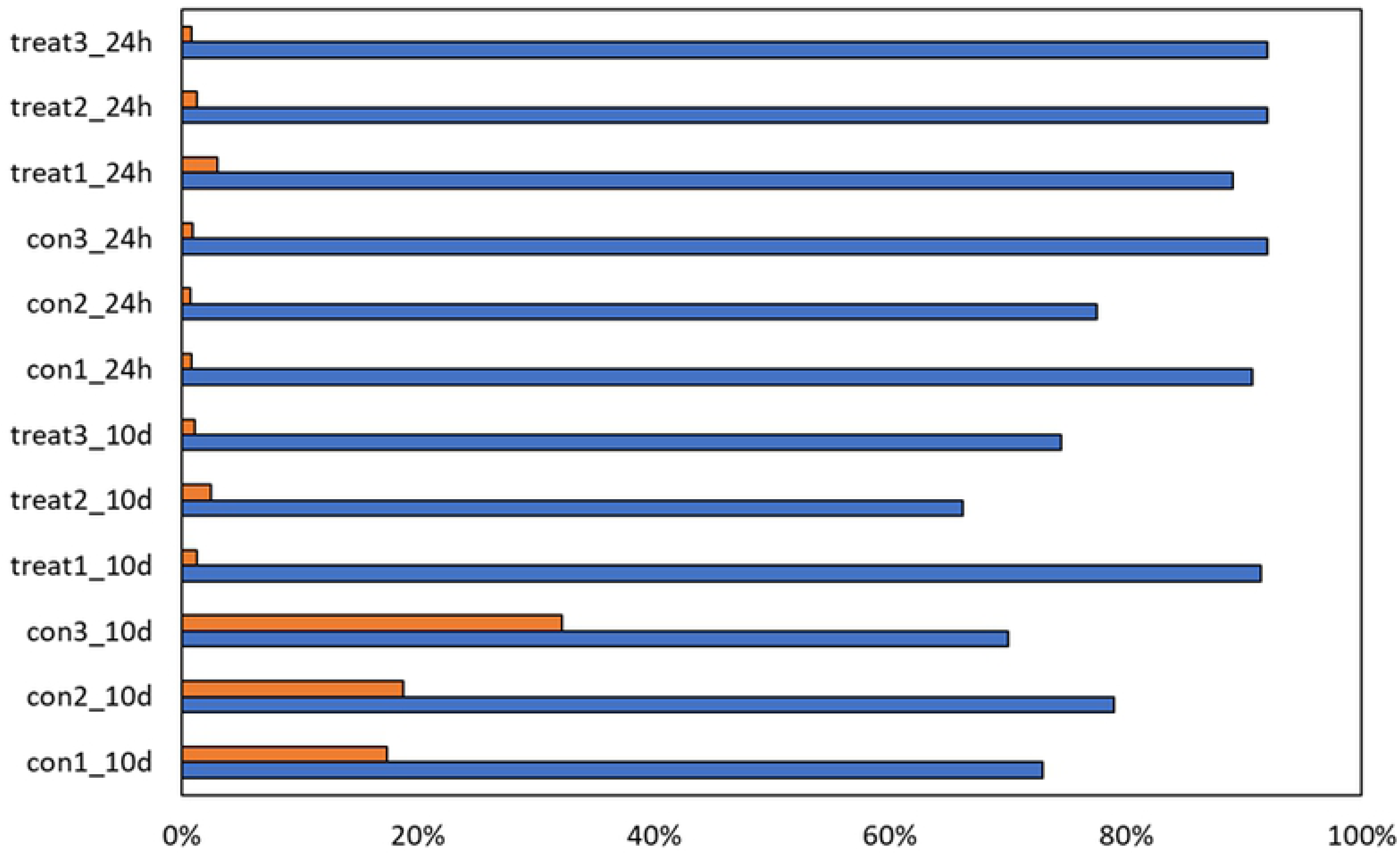
Mapped reads: Reads (%) mapped on *V. vinifera* genome (blue); reads (%) mapped on *P. viticola* genome (orange).

Principal component analysis on normalised gene expression showed a good correlation of biological replicates with the two first Principal Components (PC), explaining 90 % of variance between samples (Fig 4.). PC1, explaining 80.33 % of the variability in gene expression, separated the EO treatment from the control of the 24h experiment, whereas the second PC accounting only for a 9.47 % variation, separated the EO treatment from the control of the 10d samples. This indicates that EO induced a more important transcriptomic programming during the first 24h of vapour exposure than after the 10d treatment.

**Figure 4:**
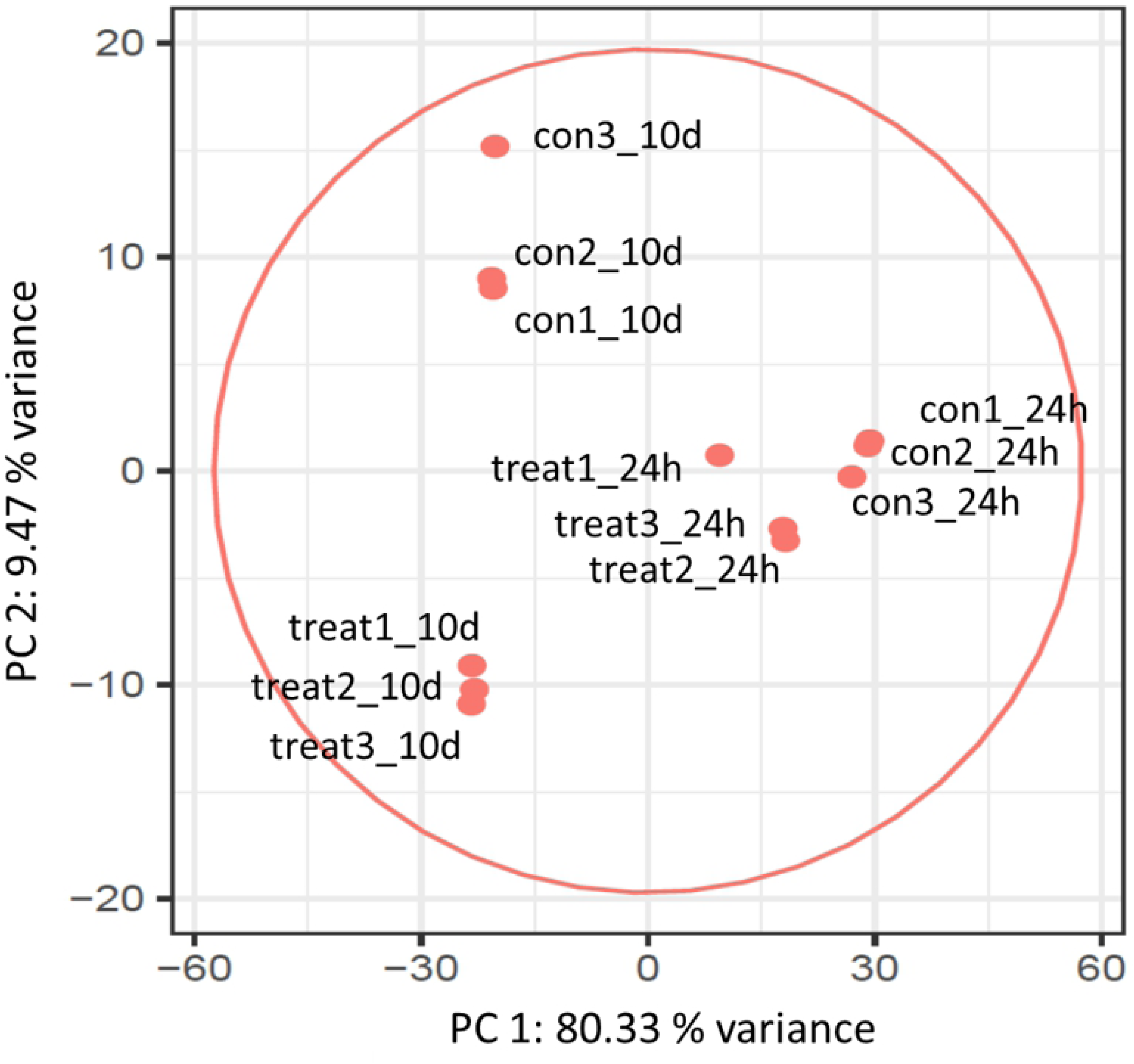
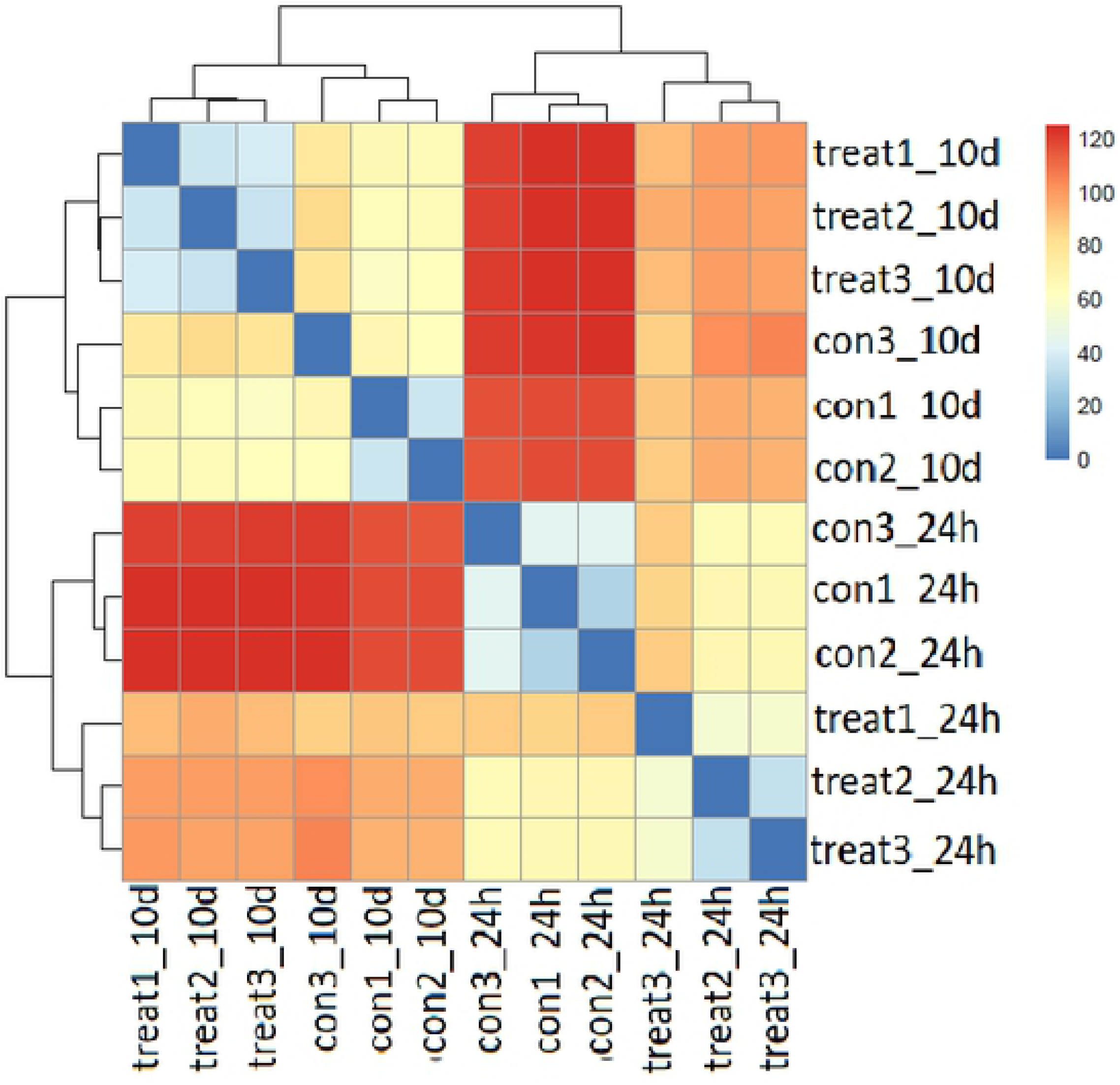
Principal component analysis (PCA) of normalised mapped reads of the control and treatments for the 24h and the 10d experiment (3 replicates for each control and each treatment).

DEG analysis of mapped transcripts on the *V. vinifera* genome yielded a total of 4800 DEGs for EO treatments after 24h and 10d. For the 24h treatment, 1061 DEGs were induced and 1189 transcripts were repressed by *O. vulgare* EO, whereas for the 10d treatment 1210 DEGs were upregulated and 807 DEGs were downregulated, respectively (figure 5). Interestingly, the number of concomitantly deregulated genes at 24h and 10d was very low. Only 40 genes were downregulated, while 37 genes were upregulated, being common to 24h and 10d treatments. In the meantime, respectively 63 and 37 genes were inversely expressed between the 24h and 10d treatments. This highlights a brief early response of the plant to EO vapour, which is only maintained during a limited time span (0 to > 24h after treatment) and is then followed by an adaptation period, where many early elicited genes return to the pre-treatment expression and a somehow long-term adaptation takes over. This early transcriptomic reprogramming conditioned by EO vapour might thus base its main effect on an eventual plant innate immunity priming.

**Figure 5:**
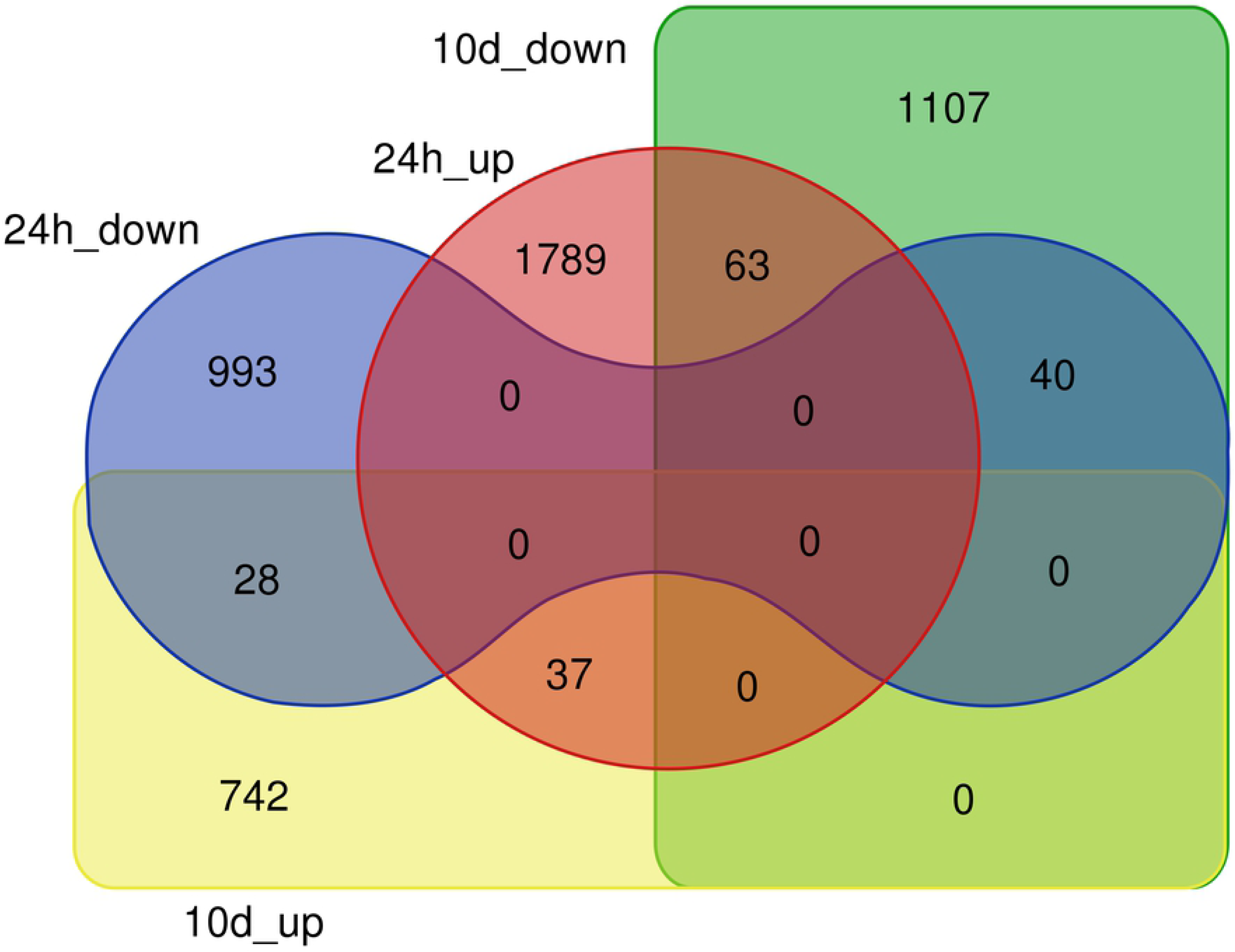
Venn diagram of differentially expressed genes (DEGs, pval>0.01, lfc<0.5) upon *O. vulgare* vapour treatment: 24h_up and 24h_down: up- respectively downregulated genes after 24h treatment; 10d_up and 10d_down: up- respectively downregulated genes after 10d treatment.

This is in some way confirmed by analysing the enriched functional categories (FC) within DEGs (Fig 6A and 6B), where for the 10d treatment, no significant enriched FC could be detected. This indicates that the long-term (10d) treatment did trigger genes that are not, or less, collectively regulated within the main metabolic pathways. It would thus be likely that the observed gene deregulation by EO in the 10d treatment is somehow random. However, for the 24h treatment several categories were significantly and highly enriched in upregulated genes (Fig 6A) as well as in downregulated genes (Fig 6B). This observation is discussed in subsequent sections below.

**Figure 6A and B:**
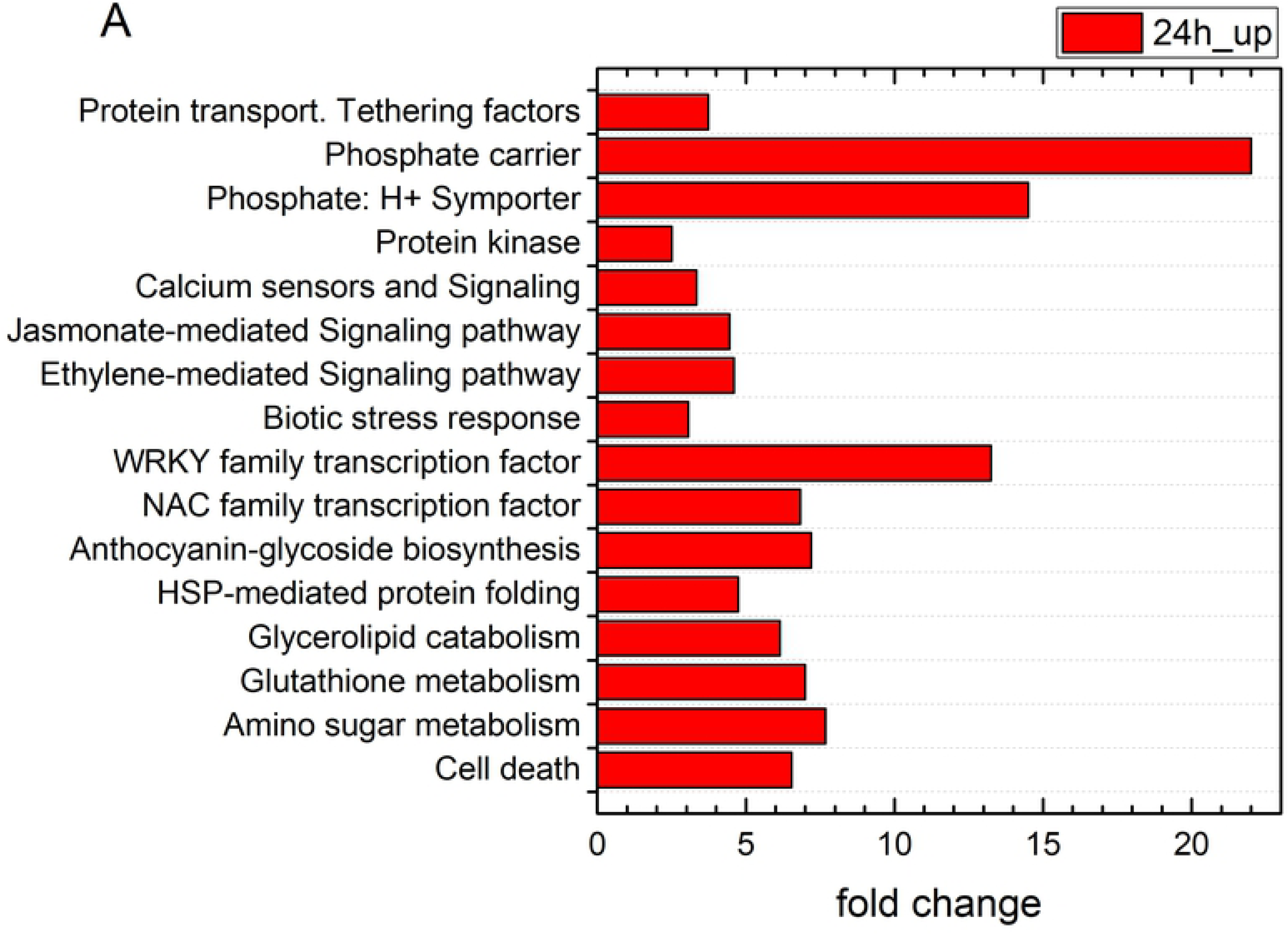
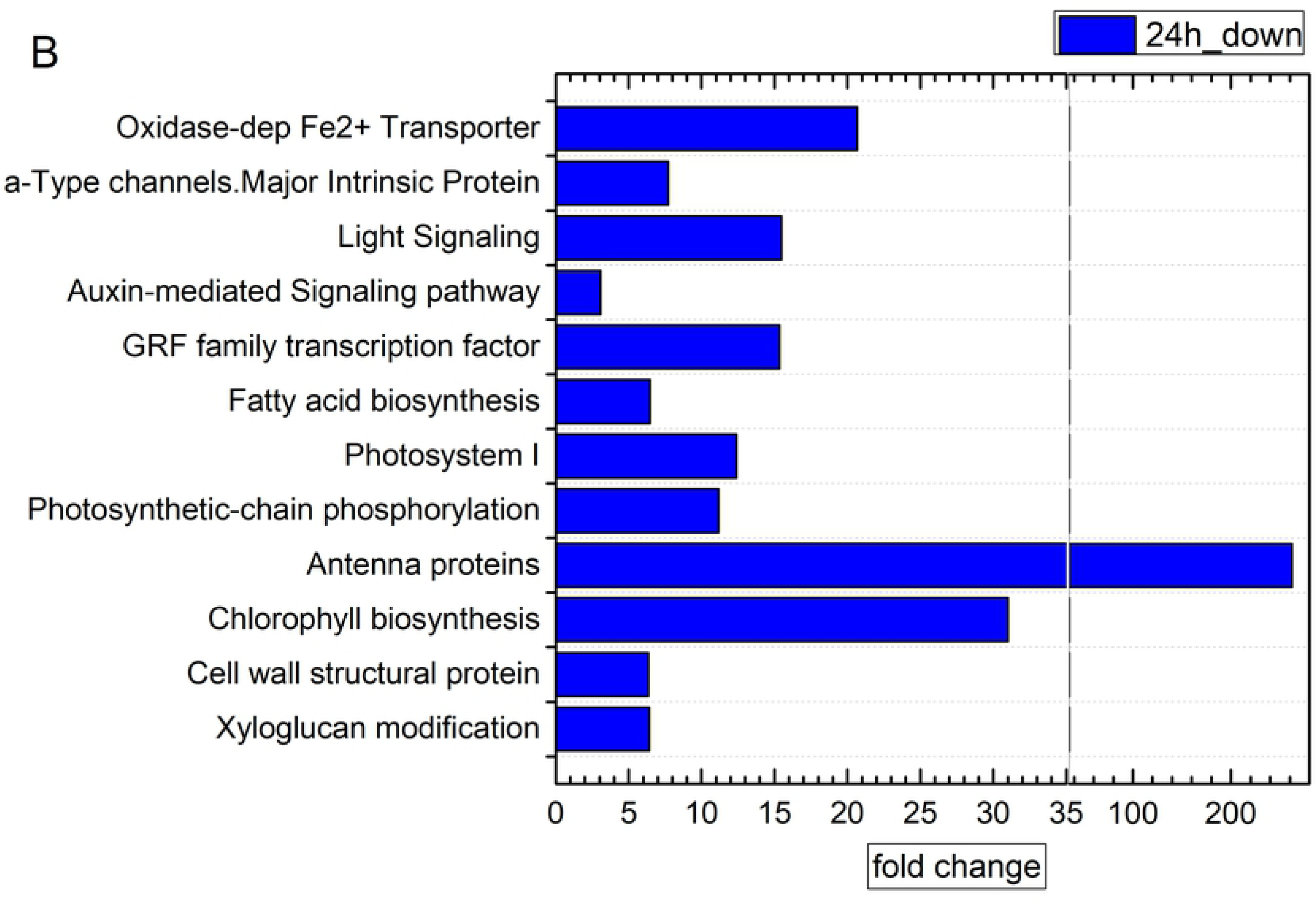
Significantly enriched functional categories in A) upregulated and B) downregulated genes in the 24h treatment. (For the 10d treatment, no functional categories were found to be significantly enriched).

Hierarchical clustering of DEGs, as illustrated by the heatmaps in Fig 7A and 7B, showed the higher number of upregulated genes for each treatment, in both 24h and 10d experiment conditions. The stringent grouping of genes detected in biological triplicates highlighted the rightness and strong significance of these results. Functional enrichment analysis of genes within each cluster, shown in Fig 8, confirmed the results obtained on total up- and downregulated transcripts as discussed above, since no significant FCs could be detected within the 10d clusters.

**Figure 7A and B:**
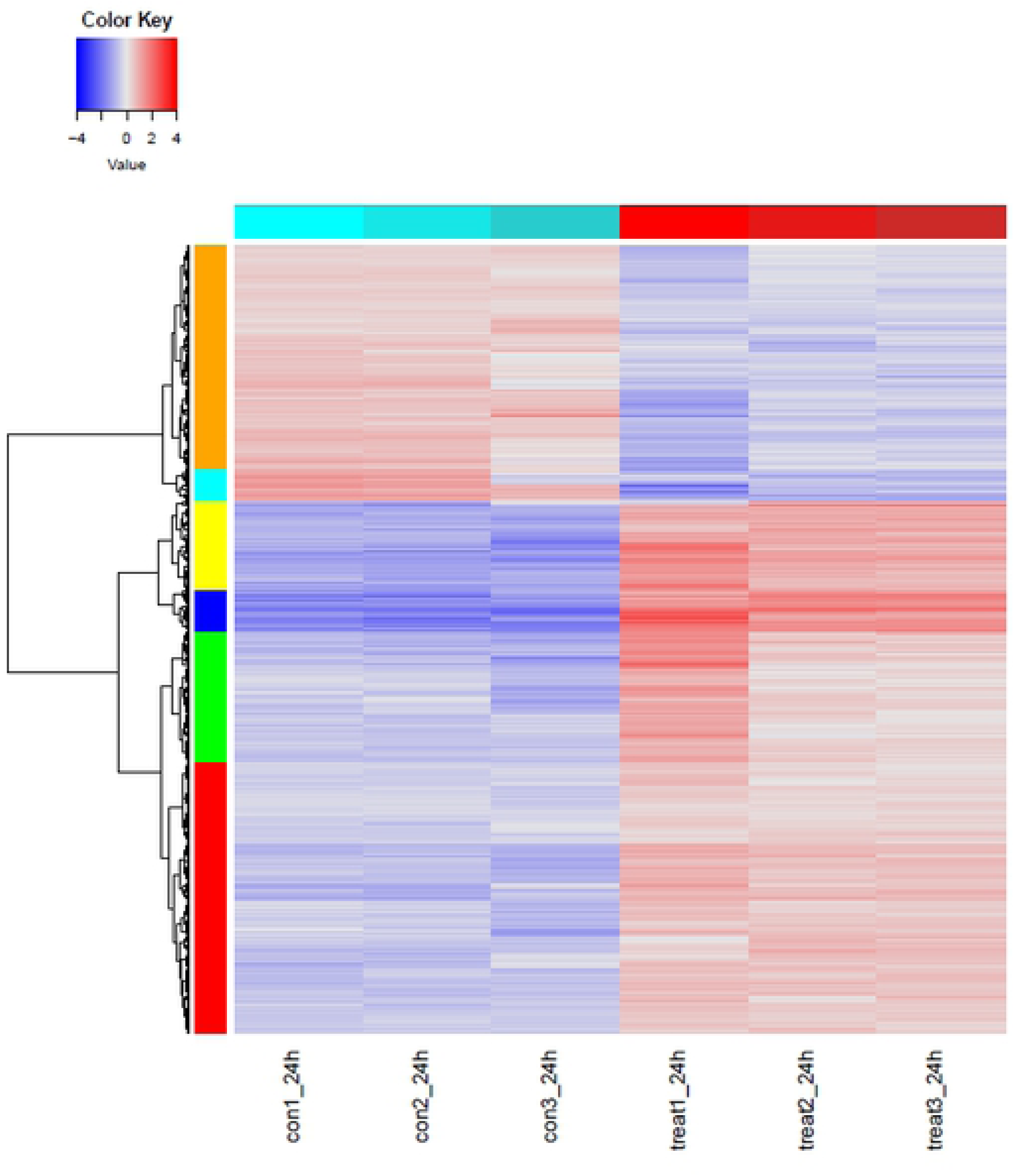
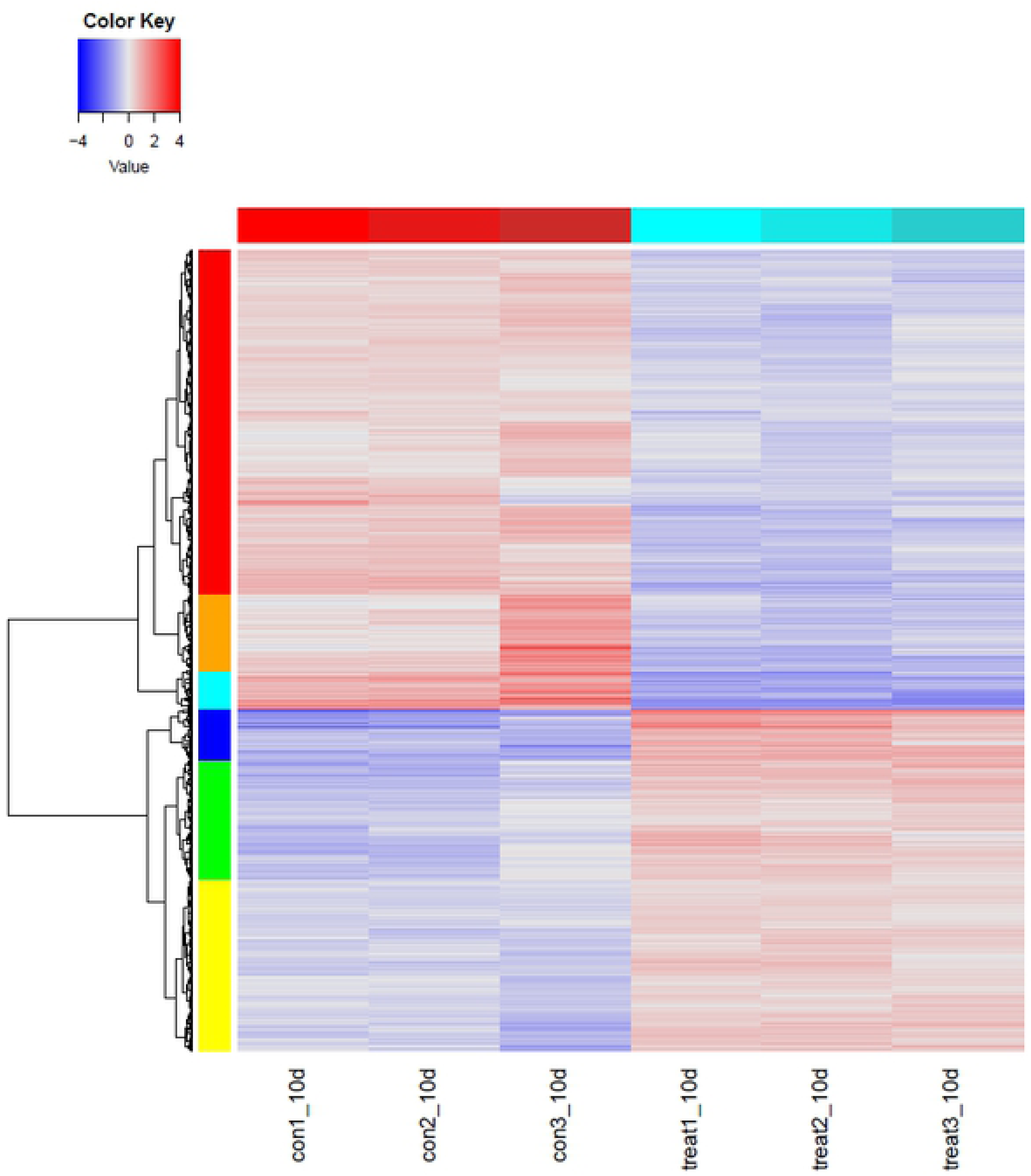
Hierarchical clustering of differentially expressed genes after A) 24h treatment and B) 10d treatment with *O. vulgare* vapour.

**Figure 8:**
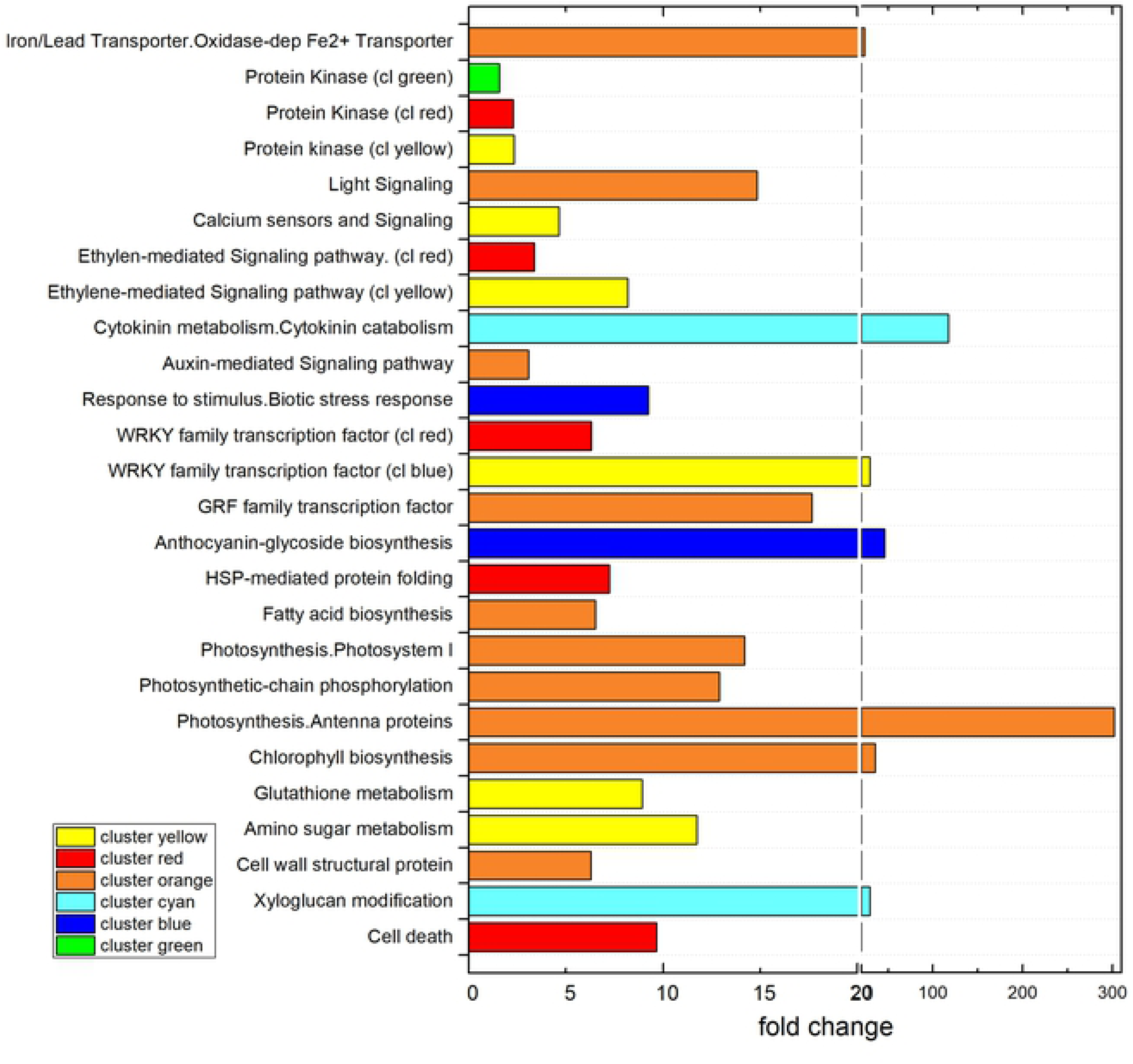
Enriched functional categories of hierarchical clusters after 24h treatment (ref. Figure 7A). (For the 10d treatment, no functional categories were found to be significantly enriched).

### *O. vulgare* vapour triggers innate plant defense mechanism controlled by hormonal signaling leading to the stimulation of phenylpropanoids

It is well established that plant responses to biotic and abiotic stress stimuli are mediated by defense-related phytohormones such as salicylic acid (SA), jasmonic acid (JA) and ethylene (ET), which act as primary signals in the regulation of plant defense (17). However, the crosstalk of hormones and downstream-induced responses are still far from being completely elucidated (46). The current state of knowledge is that SA-signaling mediates resistance to biotroph and hemi-biotroph pathogens, whereas JA- and ET-related pathways activate resistance against necrotrophs, with SA-JA crosstalk being the backbone of the plant’s immune signaling network (47).

The overview of enriched functional categories of upregulated genes, given in Fig 6A and 6B, unambiguously indicated that the innate plant immune system was primed by 24h EO vapour treatment with several highly activated metabolic pathways linked to the above-mentioned hormonal defense regulation (JA- and ET-mediated signaling), biotic stress response and secondary metabolism related to phenolic compound synthesis.

### Jasmonic acid

Amongst defense-related phytochromes, JA is thought to play an essential role in response to tissue wounding, by regulating gene expression to redirect metabolism towards producing defense molecules and repairing damage (48).

JA biosynthesis starts with the release of α-linolenic acid from galacto- and phospholipids localised on the chloroplast membrane by the action of phospholipases, which are subsequently oxidised by *Lipoxygenase* (LOX) leading to 13-hydroperoxy-9,11,15-octadecatrienoicacid (13HPOT). Two different enzyme families, termed *Allene Oxyde Synthase* (AOS) and *Allene Oxyde Cylase* (AOC), successively convert 13-HPOT into the stable cis(+)-oxophytodienoic acid (cis-OPDA) intermediate. This latter is reduced by an *Opda Reductase* (OPR) and then undergoes three rounds of β-oxidation by *Acyl-CoA Oxidase (ACX)* enzymes leading to the production of JA (49).

We showed here that EO vapour treatment triggered the expression of 3 key enzymes of the JA pathway, with *LOX* (VIT_14s0128g007901), being the key enzyme, followed by *AOX* (VIT_18s0001g11630) and *ACX3* (VIT_12s0028g02660). This points out that JA biosynthesis was highly induced by the EO treatment. Other studies reported a similar induction of *LOX* leading to a stimulation of induced resistance against *P. viticola* upon thiamin treatment of grapevines (20, 50).

Several downstream regulated genes of the JA pathway, putatively mediated by JA-signaling, were highly upregulated in EO treated plants. Amongst these latter, several isogenes of *Enhanced Disease Susceptibility (EDS1; VIT_17s0000g07400, VIT_17s0000g07420, VIT_17s0000g07370)* also known as Nonexpressor of PR Genes 1 (NPR*1*), or *Non-inducible Immunity 1 (NIM1)* or *Salicylic Acid Insensitive 1 (SAI1)*, which is an important redox sensitive transcriptional regulator of SA response and a mediator of SA-JA crosstalk (51). This *NPR1* gene is thought to be a central immune regulator for systemic-acquired resistance (SAR) and is considered as crucial for the regulation of *Pathogenesis Related (*PR) genes expression (51–53).

Consequently, several *PR Proteins 1 (PR1)* transcripts *(VIT_03s0088g00700, VIT_03s0088g00750, VIT_03s0088g00710)*, as well as several members of *the JASMONATE-ZIM DOMAIN (JAZ)* protein family *(VIT_10s0003g03790, VIT_10s0003g03800, VIT_11s0016g00710, VIT_09s0002g00890)* were induced. This latter JAZ family has been recently discovered and characterised (54) as a family of proteins orchestrating the crosstalk between JA and other hormone-signaling pathways such as ET, SA, gibberellin, and auxin. JAZs are transcriptional regulators that target different *bHLH* TFs, such as *MYCs* (55), which appeared as upregulated in the present data, too (*VIT_02s0012g01320*). This MYC branch of the JA-signaling pathway is typically activated upon wounding or feeding by herbivorous insects. The latter branch is furthermore thought to be antagonistically regulated to the ethylene-responsive transcription factor *(ERF)* branch (56). However, this has not been confirmed by the present study since *ERF*s are also highly activated as discussed the ET section. Other important regulators of the interaction between the SA and JA pathways have been identified and belong to the *WRKY* and *TGA* TF families (57). In *O. vulgare* EO treated plants a very high number of *WRKYs* such as *WRKY 6; 11; 23; 33; 40; 47;48; 51; 53; 55; 65; 70; 72* and *75*, were concomitantly upregulated with 2 *WRKY33* isogenes *(VIT_08s0058g00690 and VIT_06s0004g07500)* being amongst the highest upregulated ones (28 and 11 fold, respectively).

In general, *WRKY* TFs are a large family of regulatory proteins that are involved in various plant processes, but most notably in coping with diverse biotic and abiotic stresses. *WRKY33* functions as a positive regulator of resistance toward the necrotrophic fungi such as *Alternaria brassicicola* and *Botrytis cinerea*, (58) and was found to be induced by EO treatment in the present study.

In *Arabidopsis thaliana* L., *WRKY70* (2 upregulated isogenes: *VIT_08s0058g01390, VIT_13s0067g03140)* acts at a convergence point determining the balance between SA- and JA-dependent defense pathways while required for R-gene mediated resistance (59) and, together with *WRKY53* (*VIT_02s0025g01280*), positively modulate SAR (60). Moreover, SA biosynthesis and expression of *NPR1* also appear to be regulated by *WRKY* TFs (61). Whether *WRKY70* is indispensable for JA- and SA-signaling has, however, recently been questioned (61). This has not been confirmed by the present data, where both signaling pathways are highly activated concomitantly with *WRK70*.

### Salicylic acid

SA biosynthesis is triggered during PTI and ETI upon recognition of PAMPs or effectors of pathogens. SA is a phenolic compound that can be synthesised from the primary metabolite chorismate via two distinct enzymatic pathways, one involving *Phenylalanine Ammonia Lyase (PAL)* and the other *Isochorismate Synthase (ICS/SID2*). Diverse *PAL* coding isogenes (*VIT_16s0039g01280, VIT_16s0039g01120, VIT_16s0039g01240, VIT_16s0039g01100, VIT_11s0016g01520, VIT_11s0016g01660, VIT_11s0016g01640, VIT_16s0039g01170)* were found amongst the highest induced genes upon EO treatment. Other studies showed that a high induction of the phenylpropanoid pathway and downstream reactions also triggers, besides SA production, the production of many phytoalexins. For example, a high *PAL* activation, leading to an enhanced phenolic compound accumulation, provided a fair protection against leaf stripe caused by *Drechlsera graminea*, as reported on barley treated with aqueous leaf extract of *Azadirachta indica Juss*. (62).

Besides enzymes that directly contribute to SA biosynthesis (*ICS* and *PAL*), other proteins have been identified as participating in pathogen-induced SA accumulation. These include *EDS*1/*NPR1* (as already discussed above for JA) and *PAD4 (Phytoalexin Deficient 4; VIT_07s0031g02390*), highly upregulated by EO vapour (63). Similar results were observed when *A. thaliana* was treated with *Gaultheria procumbens* L. EO, inducing an SA-mediated defense response and resistance to *Colletotrichum higginsianumin* Sacc. (64). However, these authors demonstrate that this effect is mainly due to the methyl salicylate present in the used EO, which triggered the SA-mediated defense response. Another microarray study was carried out on *A. thaliana* plants treated with different commercial herbal plant preparations, and showed that SA- and JA defense responses could be induced together (65). As far as we know, the induction of an SA-JA-mediated defense response triggered by the vapour phase of *O. vulgare* has never been reported previously.

Other SA-related enzymes, which play an important role in plant-pathogen interactions are *Glutathione S-Transferases (GSTs)*. In the present study, several tau class *GSTs*, GSTUs (GSTU 1; 8; 9; 10; 20; 22) and a very high number of *GSTU 25* isogenes were the most upregulated transcripts, particularly a *VIT_19s0015g02730*, a *GSTU 25*, which were induced more than 128 times. Elevated *GST* activities have often been observed in plants treated with such beneficial microbes (bacteria and fungi), resulting in the induction of a systemic resistance response to subsequent pathogen infections. Their confirmed roles in biotic and abiotic stress tolerance are due to a detoxification capacity by conjugation with glutathione, the attenuation of oxidative damage and their contribution in hormone transport. The exact metabolic functions and their contribution to disease resistance in plants, however, remain to be elucidated (66).

### Ethylene and Ethylene-mediated signaling

ET is a principal and pleiotropic modulator of many aspects of plant life, including various mechanisms by which plants react to pathogen attacks. ET leads to a cascade of transcription factors (TF) consisting of primary *Ethylene-insensitiv (EIN)* 3-like regulators and downstream *ERF* (*Ethylene Response Factor)*-like TF, controlling the expression of various effector genes involved in various aspects of systemic-induced defense responses. Moreover, as discussed above, a significant cross-talk occurs with other defense response pathways controlled by SA and JA, eventually resulting in a differentiated response to disease that is not very well understood so far.

ET biosynthesis involves the conversion of S-adenosyl-Met (SAM) to 1-aminocyclopropane-1-carboxylate (ACC) and methylthioadenosine, performed by the enzyme *ACC synthase (ACS)* and a further step where ACC is converted to ET, CO_2_, and cyanide by *1-aminocyclopropane-1-carboxylate oxidase (ACO)*, whereas conversion of AdoMet to *ACC* by *ACS* is generally considered the rate-limiting step (46).

In our study, upon *O. vulgare* EO treatment, several *ACOs (VIT_02s0012g00400, VIT_05s0049g00410, VIT_01s0011g05650, VIT_12s0059g01380)* and one *ACS 1 (VIT_02s0025g00360)* were induced, indicating that ET synthesis was activated by the EO treatment. Interestingly, a recently discovered TF, *MBF1c (VIT_11s0016g04080)*, which acts upstream of ET and SA, and was thought to be only involved in abiotic stress responses such as heat stress in *A. thaliana* (67, 68) and grapevine (69–71), was found to be upregulated here. Accordingly, a cascade of ethylene-controlled *ERFs* was highly upregulated, notably an *Apetala2 gene (AP2)/ERF (VIT_11s0016g00660)*, which was overall the second most upregulated transcript in the 24h treatment. The *AP2/ERF* superfamily plays a pivotal role in adaptation to biotic stresses (72), emphasising that the innate immune response of the plant is highly active. This was confirmed by other highly induced *ERFs*, notably an *ERF109 (VIT_03s0063g00460)*, which plays an important role in abiotic stress adaptation, as reported for salt tolerance (73). Two other ERFs related to *AP2 (VIT_18s0072g00260* and *VIT_11s0016g00670)* were simultaneously upregulated up to 50 times by the EO treatment. Interestingly, *ERF 105 VIT_16s0013g00980* seems to play a complex role in biotic and abiotic regulation in different tissues, just as it has been shown, for example, to be highly heat stress responsive (71), as well as circadianly regulated in grape berry tissue (70).

In general, *ERF*s act as a key regulatory hub and integrate ET, ABA, JA, and redox signaling in the plant response to a number of abiotic and biotic stresses, such as those caused by pathogens, wounding, cold and heat stress, UV light, drought, and salinity (72). Their upregulation in the present experiment have given an indication of their role within the regulation of the innate immune system upon exogenous elicitors, and pointed out their important role within plant defense mechanism to biotic stresses. The set of target genes regulated by each ERF has not yet been completely elucidated, and the present gene expression data might contribute to the understanding of complex coregulations. ERFs activate the transcription of basic type defense-related genes, such as pathogenesis-related (PR) genes, *osmotins, chitinase and β-1,3-glucanase genes*. This also happened in the present study where several PRs putatively controlled by ERF, such as *β-1,3-glucanases (VIT_06s0061g00120, VIT_08s0007g06060, VIT_08s0007g06040), chitinases (VIT_05s0094g00220, VIT_16s0050g02230)* and *osmotins (VIT_02s0025g04280, VIT_02s0025g04250*), were simultaneously upregulated.

These latter are grouped in different PR families. Chitinases, belonging to the PR-3 protein group, are hydrolytic enzymes that break down glyosidic bonds in chitin, which is a major component of the cell wall of pathogenic fungi. Chitinases make the fungi inactive without any negative impact on the plants and can thus enhance the plant’s defense system (74). The fact that *P. viticola* is an oomycete and, as such, does not contain chitin, highlights that the triggered plant immune response is unspecific to the pathogen and might thereby impede infection with other pathogens as well. The *β-1,3-glucanases* are grouped in the *PR-2* family of PR proteins and are involved in plant defense by hydrolysing the cell walls of fungal pathogens most commonly in combination with chitinases. *In vitro* analysis has shown that b-1,3-glucanases directly act on fungal pathogens by degrading *b-1,3/ 1,6-glucans* and that chitinases act by attacking the bond between the C1 and C4 of two consecutive N-acetylglucosamines of chitins in the fungal cell wall (75).

It was shown that they also indirectly act as defense mechanisms elicitors by releasing b-1,3-glucan and chitin oligosaccharides (75). Glucans have been shown to play a major role in grapevine immunity against *P. viticola* and could be elicited by treatment with sulfated laminarin (76). We here show the possibility that these defense PRs could be triggered by EO treatments.

*Osmotins* are *PR-5s* proteins and have been shown to play an important role in response to biotic and abiotic stresses in plants, which role is mainly activated by *MAPK (Mitogen-activated protein kinases)* pathways. Osmotin proteins enter the fungal plasma membrane and activate the defense system. They are also involved in the initiation of apoptosis and PCD (75).

Their expression was shown to be induced by a number of different signals as, for instance, by SA, ABA, auxin, ET, salinity, drought conditions, UV light, wounding, desiccation, cold, fungal infection, oomycetes, bacteria and viruses. The present data suggests that osmotin synthesis would also be triggered by *O. vulgare* vapour which, as far as we know, has never been reported in literature before.

### Mitogen-activated protein kinases (MAPKs) and CA-signaling induction

MAPKs are important regulators of plant immunity that generally transduce extracellular stimuli into cellular responses. These stimuli include the perception of PAMPs by host transmembrane pattern recognition receptors, which leads to PTI. In the A. thaliana model, molecular genetic evidence implicates a number of MAPK cascade components in PAMP signaling, and in responses to immunity-related phytohormones such as ET and SA (77). In Arabidopsis, MPK3, MPK6, MPK4, and MPK11 are rapidly activated during PTI (78) but also in ETI, where MPK3/MPK6 seem to play an essential role (79).

In response to *O. vulgare* EO treatment, several MAPK cascade members, such as a *MPK3 (VIT_06s0004g03540)* and a *MPK13 (VIT_06s0004g03620*) were upregulated. Shoresh et al. (2006) showed that *MPK3* orthologues conferred immunity to cucumber plants against *Pseudomonas syringae pv. lachrymans* (80) and their overexpression created autoimmune phenotypes characterised by dwarfism associated with spontaneous cell death, accumulations of ROS and SA, and modification of phytoalexin metabolism (81).

In addition to MAPK cascade signaling, PAMP perception was shown to induce Ca^2+^ dependent kinases (CDPKs) by regulating Ca^2+^ influx channels (82). Recent findings indicate that Ca^2+^- ATPases regulate Ca^2+^ efflux as well as innate immune defenses (83, 84). Our results pointed in the same direction with an upregulation of FCs related to calcium sensors and Ca-signaling (Fig 6.) upon *O. vulgare* EO treatment. Within these categories several *Calmodulin* isogenes were found to be highly overexpressed, as well as *Ca2+-ATPases (VIT_05s0020g04300, VIT_07s0129g00180, VIT_07s0129g00110)* and several transcripts, coding for Calcium-binding proteins *(VIT_01s0010g02950, VIT_08s0056g00290 VIT_01s0026g02590)*.

Hypersensitive Response (HR) is associated with the innate immune response of plants and commonly regarded as a feature of ETI. This particular response involves programmed cell death (PCD) and occurs at the point of pathogen entry, resulting in an efficient containment of the pathogen (85). Reactive Oxygen Species (ROS) have been well established as an integral aspect of plant immunity in a process generally described as the oxidative burst involved in PCD and thus in HR (86). In the present study, the apoptosis associated FC was highly enriched in EO-triggered genes, pointing out that this part of the innate immune response was activated along with a *Respiratory Burst Oxidase Protein F (RBOHF)* transcripts (*VIT_02s0025g00510*), known to be regulated by abscisic acid (ABA), whose biosynthetic enzyme *9-cis-epoxycarotenoid dioxygenase* (NCED) *VIT_19s0093g00550* was induced as well. This highlighted a contradiction sometimes encountered in transcriptomic literature, where JA/ET are often reported to be antagonistically regulated, notably in abiotic stress studies (87). Other genes related to *ROS* production, such as *dicyanin, dicyanin blue copper protein, glutaredoxin*, as well as several *peroxidases* and an *alternative oxidase* (*VIT_02s0033g01380*, 18 fold upregulation), which are crucial in the early generation of mitochondrial ROS and precede fungal elicitor-induced HR or a fungal toxin-induced cell death (88), were found upregulated, which highlighted that the oxidative stress response was triggered by EO vapour. This was concomitant with the upregulation of several Heat Shock TFs, which can function as molecular sensors that directly sense ROS and control the expression of oxidative stress response genes during oxidative stress (Miller and Mittler, 2006).

### Phenylpropanoid synthesis is highly activated by *O.vulgaris* vapour

Phytoalexins are defined as low-weight antimicrobial secondary metabolites that are synthesised and accumulate in plants after pathogen attacks, but can be activated by PTI (89) as well.

Many reported phytoalexins represent phenylalanine-derived phenylpropanoids. As previously discussed, this latter pathway regulated by the key enzyme *PAL* was highly activated upon EO treatment. Downstream of *PAL*, the rate-limiting enzyme of flavonoid/isoflavononid biosynthesis, *chalcone synthase (CHS; VIT_16s0100g00860)*, was highly induced as well, indicating an enhanced synthesis of those phenolic compounds. Besides being part of the plant developmental programme, the *CHS* gene expression was reported to be induced in plants under stress conditions such as UV light, bacterial or fungal infection. *CHS* expression causes accumulation of flavonoid and isoflavonoid phytoalexins and is involved in the SA defense pathway (89, 90). Respectively a *flavanol synthase* coding transcript *(VIT_18s0001g03430)*, as well as a *Quercetin 3-O-methyltransferase (VIT_02s0025g02920)*, which were amongst the top ten induced genes, two *Leucoanthocyanidin dioxygenase (LAR)* coding isogenes *(VIT_13s0067g01020; VIT_11s0118g00360)* and several genes related to Anthocyanin biosynthesis, were concomitantly induced by the EO treatment. Studies on cucumbers showed a similar increase of *PAL* activity and other genes of the flavonoid pathway, after application of Milsana®, a resistance inducer, also known for its direct antimicrobial effect (91).

Amongst the most important representative of phenolpropanoids-derived phytoalexins are stilbenes, notably resveratrol, which accumulates following infection in several phylogenetically unrelated species, including grapevine (92). The biosynthesis of this secondary metabolite requires the presence of only one unique enzyme, *stilbene synthase (STS)*, which has thus become a feasible molecule for metabolic engineering. For example, heterologous expression of the grapevine *STS* gene *VST1* under control of its native promoter in tobacco (*Nicotiana tabacum*) has led to pathogen-inducible biosynthesis of resveratrol, which correlated with enhanced disease resistance to *B. cinerea*. Upon *O. vulgare* treatment, 9 isogenes coding for *STSs* were upregulated to very high levels such as *VIT_10s0042g00890* and *VIT_10s0042g00870* with a respectively 42- and 32-fold induction. In addition, several *Myb14 TF* coding isogenes (*VIT_07s0005g03340, VIT_12s0134g00480, VIT_07s0005g03340*) were highly upregulated (20 to 22 fold). The latter is, together with *MYB15*, the transcriptional regulator of stilbene biosynthesis in grapevine (93). Enhanced accumulation of stilbenic phytoalexins has been shown to be implicated in the resistance of grapevine cultivars to three major fungal pathogens, *B. cinerea, P. viticola* and *Erysiphe necator* (powdery mildew) (10), and could thus be amongst the main reasons why *P. viticola* development was inhibited under *O. vulgare* treatment. An overview of deregulated transcripts within the phenylpropanoid pathway is given as a supplemental in figure S5.

Stimulation of the phenylpropanoid pathway and notably stilbene synthesis is highly regulated by ET and JA, as shown by exogenous application of JA or ET in several studies (94–96), which also seems to be the case in the present experiment.

### *O. vulgare* impedes the photosynthetic machinery and cell wall synthesis related processes

Visual observation as well as photosynthesis (PS) measurements (Fig 2) during EO vaporisation showed a phytotoxic effect of EOs on vine physiology, which correlated with gene expression data.

Several FC related to PS are enriched in repressed genes, as shown in Fig 6B (antenna proteins, light signaling, PSI, PS chain-reaction, Chlorophyll biosynthesis). Within these categories, 18 light-harvesting complex (LHCs) coding transcripts were repressed by EO treatment after 24h. Interestingly, some were no longer repressed or even induced after a 10d treatment, as is the case of *LHCB6 (VIT_12s0055g01110)*. A similar observation can be made regarding genes involved in PSI (Photosystem I reaction center subunit genes), and PSII genes linked to photosynthetic-chain phosphorylation and chlorophyll biosynthesis, that were systematically repressed after a 24h treatment but no longer after 10d. This would indicate that the primary reactions transiently disappear with an increased duration of treatment and suggest some kind of adaptation to EO vapour.

A direct phytotoxic effect of EO has previously been reported in several studies (41) and mainly attributed to carvacrol and thymol (97–99), which are also the main terpenes of the applied EO in the present study.

However, downregulation of PS and PS-related processes could also be part of the innate immune plant response, which has been described in several studies. In particular, upregulation of MPK6, as discussed above, has been associated with an altered expression of photosynthesis-related genes and inhibition of photosynthesis (79).

### Conclusion

The need for sustainable alternatives to replace or reduce the use of synthetic pesticides in agriculture is of utmost urgency. EOs that were shown to possess antifungal properties in many previous studies, could potentially represent a natural strategy to replace or at least reduce the use of synthetic fungicides. However, their adhesiveness and stability on the plant is very bad, when applied in their liquid phase. The present study is, as far as we know, the first one to test and confirm that the vapour phase of EOs is *in vivo* highly efficient against the major pathogen *P. viticola*, the causing agent of downy mildew. This would offer new alternatives in the development of innovative plant protection strategies that would involve fumigation systems in greenhouses or dispensers in field-grown crops, using volatile organic compounds from plant EOs.

Even more significantly, the present study provides important information regarding the underlying mechanisms in the host-pathogen-EO interaction and shows clearly that the *O. vulgare* vapour triggered a multilayered immune system of the plants. Gene expression analysis revealed a complex activation of hormonal crosstalk involving JA, ET, and SA biosynthesis and their signaling cascades. This led to the activation of different immune mechanisms involving PR genes activation, flavonoids and stilbene synthesis, as well as transcriptional networks in relation to PCD and apoptosis. Not only does *O. vulgare* vapour, or one or several of its constituents, seem to act as PAMPs to trigger PTI, but also as an elicitor that transiently triggers ETI. This complex interaction is confirmed by numerous studies that have shown that ETI, basal defense and PTI use a common set of signaling components including multiple regulatory proteins, reactive oxygen intermediates (ROIs), as well as the phytohormones salicylic acid (SA), ethylene (ET) and jasmonic acid (17).

To what extent the inhibitory effect of EO vapour on *P*. v*iticola* development was due to a direct toxicity of the EO vapour on the pathogens or to the stimulation of the innate immune system of vines, could not be clearly elucidated with the applied methodology.

The study, however, provides important molecular data, such as target genes, gene networks and metabolic pathways involved in the innate immune system of the plant and therefore important for future genetics studies and resistance-breeding programmes (100–102).

## Material and Methods

### Experimental set up

Two customised vaporisation systems (control and treatment) were built to enable a continuous flow of EO vapour during several weeks. Inside a basic climatic chamber (CLF Plant Climatics, Model L-66LL VL), an air-tight Plexiglas chamber was fitted top and bottom with connections for the vaporisation hose (supplemental figure 1). EOs were put in a Petri dish, inside two plastic boxes, which could optionally be heated to 35°C. The two Plexiglas chambers (control and treatment) were connected to a continuously running custom-made compressor. This way, vapour was distributed from the plastic boxes containing Petri dishes, with or without EO, inside the climatic chamber by the bottom tube and extracted by the exhaustion tube connected to the top of the chamber. Plants were put on an alveoled platform to ensure a homogenous distribution of vapour inside the chamber.

Because, according to literature, the most efficient EOs against bacteria and fungi come from the species belonging to the *Origanum* and *Thymus* genera (28, 38), these were chosen to be tested during the experiment

Commercially available standard essential oils were purchased from Compagnie des sense^®^ (France). Components of these EOs were analysed by a gas chromatography (Agilent 7890B; Capillary column: 60m, 0.25mm ID, 1.4µm, Rtx®-1301, vector gas: hydrogen, flow 4mL.min^−1^, injector temperature 100°C, Gradient: 5 min at 40°C then 3°C.min-1 until 240°C, total runtime 71.67 min) with Flame Ionization Detector (FID) (250°C, Air flow 400mL.min^−1^, H2 fuel flow 30mL.min^−1^). Composition of EOs are provided in supplemental file S2.

The EO vapour concentration inside the chamber during vaporisation was also assessed by GC-FID. For this purpose, active charcoal was placed at different spots inside the control and vapour chamber, and subsequently extracted by dichloromethane and injected in the GC-FID system.

### Plant material and experimental conditions

Each experiment was carried out with two-year-old cuttings of cv Chasselas (*V. vinifera*) at a 12 to 15 leaf stage. All leaves of the 12 plants were artificially infected with *P. viticola*, using a suspension containing 10^5^ sporangia.mL^−1^, which was sprayed on the lower site of each leaf. Inoculated plants were subsequently split into two groups of 6 plants for control and treatment, then put inside their respective climatic chamber. Infection was performed at the end of the day so as to provide optimal infection conditions for *P.viticola*, i.e. overnight in the dark and in presence of a humidifier.

The appearance of oil spots on control plants (untreated) indicated that incubation of *P. viticola* was completed. Plants were then moistened in the evening to provoke sporulation during the night and allow better visual assessment and easy identification of infected leaf tissue for RNA-seq sampling.

Two series of experiments were conducted with each EO, resulting in a total of 4 sets of 10d experiments. The two first series consisted in a continuous treatment with each EO vapour (*T. vulgaris* and *O. vulgaris*) for a duration of 10d starting straight after infection. Plants in the control chamber were infected with *P. viticola* but not treated with EO vapour, however a continuous air flow with the same debit as in the treatment was applied throughout the whole period.

For the second set of experiments, plants were also treated immediately after infection with *P. viticola*, but vaporisation was maintained for only 24h. Plants were subsequently maintained in the chamber for a total of 10d (without treatment), until oils spots eventually appeared on the control plants.

In order to assess a putative phytotoxicity effect, physiological parameters were recorded every 2 to 4 days. Leaf emergence rate (LER) as an estimate for growth speed, was assessed by leaf counts 3 times a week and photosynthesis was measured by gas exchange measurements with a Ciras 3 (PP systems), USA Environmental parameters inside the chambers were Photosynthetically Active Radiation (PAR) of 500 mmol.m^−2^s^−1^, a temperature of 26/20 °C (respectively day/night) and a relative humidity of 50 %.

### RNA extraction

Leaves that showed a maximum number of spores after induction of sporulation (assessed by visual inspection), were sampled after 10d of continuous fumigation in the first set of experiments and 24h post-inoculation coinciding with the end of the vaporisation in the second set of experiments.

RNA extraction was performed according to the protocol described by Rienth et al. (103). Briefly, one gram of leaf matter was ground to powder under liquid nitrogen. Subsequently, 5 ml of extraction buffer (6 M guanidine-hydrochloride, 0.15 M tri-sodium-citrate, 20 mM EDTA and 1.5 % CTAB) was added. Cell debris were removed by centrifugation. After chloroform extraction, one volume of isopropanol was added to the resulting aqueous phase to precipitate RNAs. Samples were kept at −20 °C for at least two hours. Precipitated RNAs were separated by centrifugation and cleaned with 75 % ethanol. The pellet was re-suspended in the RLC buffer of RNAeasy® Kit (Qiagen, Switzerland) and an additional chloroform-cleaning step was applied. The successive washing steps and the DNAse treatment were performed following the manufacturer’s recommendations. Optical densities were measured at 260 and 280 nm with a NanoDrop 2000c Spectrophotometer (Thermo Scientific, Switzerland).)

### Transcriptome sequencing

Total RNA samples were qualitatively and quantitatively controlled using a Bioanalyser 2100 (Agilent, Switzerland) and a Qubit® 3.0 Fluorometer (Thermo Fisher Scientific, Switzerland). Libraries were created with the TruSeq Stranded mRNA sample preparation kit (Illumina, Switzerland) following the manufacturer’s recommendations. The libraries’ quality was then checked with the Fragment Analyser (AATI – Agilent, Switzerland). Transcriptome-sequencing was carried out within one Illumina MiniSeq run at 2 × 151 bp paired-end read length, using a MiniSeq High Output kit (Illumina). Total sequencing yielded 12.4 Gbp and generated between 0.73 Gbp and 1.49 Gbp per sample. Reads were automatically trimmed for adaptor removal and demultiplexed using the BaseSpace Sequence Hub (Illumina). The quality of reads was performed with FastQC version 0.11.8 (http://www.bioinformatics.babraham.ac.uk/projects/fastqc). A final reads trimming (Illumina adapters removal) and filtering step was performed using Trimmomatic software version 0.36 with a specified average quality cut-off of 30 (Phred score) and a minimum read length of 40 bp.

The raw sequences reads of the twelve samples have been made publicly available as fastq files, in the Sequence Read Archive (SRA) database of the National Centre for Biotechnology Information (NCBI) (104), under the following accessions SRR8439286, SRR8439287, SRR8439284, SRR8439285, SRR8439288, SRR8439289, SRR8439290, SRR8439291, SRR8439292, SRR8439293, SRR8439294, SRR8439295 (see supplementary Table 1 for details).

### RNA sequences analysis

Cleaned reads were first mapped against predicted mRNAs (PN40024 12X v2 grape reference transcriptome) obtained from the gene prediction version 2.0 of NCBI. Reads were also mapped to the *P. viticola* genome available at DDBJ/ENA/GenBank under the accession MTPI00000000 (45), to check presence or absence of the fungi inside the leaves.

Mapping was performed using HISAT 2 (105) followed by read counting with HTSeq (106). Differential expression analysis was performed using the DEseq 2 package for R (107). A principal component analysis was performed with R on all normalised reads. Transcripts were considered as differentially expressed (DEG) when adjusted p-value was < 0.01 and log2-ratio was > 0.5 (supplemental file S4).

Hierarchical clustering by heatmap was performed with DEseq2 on log transformed DEGs. To have an overview of similarities and dissimilarities among samples, the count data were used to perform heatmap analysis with hierarchical clustering and principal component analysis (PCA) with DESeq2 R package. Venn diagrams were drawn with tools provided by the Center of Plant Systems Biology at Ghent University http://bioinformatics.psb.ugent.be/webtools/Venn/. Gene annotation was derived from Grimplet et al., 2012. Functional Categories of transcripts up- and downregulated and allocated to different clusters were analysed with FatiGO (108), to identify significant enrichment of functional category. Categories were derived from Grimplet et al (2012) (109) and Fisher’s exact test was carried out to compare the genes list with non-redundant transcripts from the grapevine genome. Significant enrichment was considered in case of p value < 0.05 and illustrated as fold change. For the illustration of the phenylpropanoid pathway, transcripts that were significantly and concomitantly modulated (fc >2, p<0.05) in either both green or both ripe stages were mapped using VitisNet networks through cytoscape v 2.8.3 s.

## Supporting information

S1: Picture of the custom-made climatic chamber and vaporisation system S2: Oil composition and concentration inside the chamber

S3: Summary tables with sequence details

S4: Differentially expressed genes with annotations and clusters

S5: Cytoscape graph of affected transcripts within the phenylpropanoid synthesis pathway. Blue: repressed, red: induced transcripts by 24h EO vapour treatment. Parallelograms: RNA; Round rectangles: proteins; Ellipse: simple molecules; Diamonds: state transitions, transcription and translation.

S6: Genes allocated to clusters of Figures 7A and B.

## Author Contributions

Conceptualization, M.R and F.L.; Ideas, M.R and F.L., Data Curation, M.R. and J.C., Formal Analysis, M.R. and F.L., Funding Acquisition, M.R. and F.L., Methodology, M.R., J.C. and F.L.; Investigation, M.R., S.G., J.C., F.L.; Project Administration, M.R. and F.L.; Resources, M.R., J.C. and F.L.; Supervision, M.R. and F.L.; Validation, M.R. and F.L.; Visualization, M.R. and F.L.; Writing - Original Draft Preparation, M.R. and F.L.; Writing - Review & Editing, M.R. and F.L.; Resources, M.R. and F.L.; Supervision, M.R. and F.L.

## Acknowledgments

The authors are grateful to the HealthFood Research Programme of the HES-SO University of Applied Sciences and Arts Western Switzerland for funding this study. The authors also gratefully acknowledge Marylin Cléroux for the GC-FID analysis, Arnaud Pernet for constructing the vaporisation system, Jean-Philippe Burdet for fruitful discussions and suggestions, Raphael Gonzales and Eric Remolif for support with plant cutting preparations and Romain Chablais for technical assistance during sequencing work. Furthermore authors would like to thank the COST (European Cooperation in Science & technology) - action CA17111 – INTEGRAPE that permitted valuable exchange with another scientist on data analysis and integration and the Swiss Government excellence scholarship for financing the postdoctoral position of Dr. Sana Ghaffari.

## References

1. Gisi U, Sierotzki HJE. Fungicide modes of action and resistance in downy mildews. Eur J Plant Pathol. 2008;122(1): 157–167.

2. Kim K-H, Kabir E, Jahan SA. Exposure to pesticides and the associated human health effects. SciTotal Environ. 2017;575: 525–535.

3. Gessler C, Pertot I, Perazzolli M. *Plasmopara viticola*: a review of knowledge on downy mildew of grapevine and effective disease management. Phytopathol Mediterr. 2011;50(1): 3–44.

4. Burruano S. The life-cycle of *Plasmopara viticola*, cause of downy mildew of vine. Mycologist. 2000;14:v179–182.

5. Furuya S, Mochizuki M, Saito S, Kobayashi H, Takayanagi T, Suzuki S. Monitoring of QoI fungicide resistance in *Plasmopara viticola* populations in Japan. Pest Manag Sci. 2010;66(11): 1268–1272.

6. Chen W-J, Delmotte F, Cervera SR, Douence L, Greif C, Corio-Costet M-F. At least two origins of fungicide resistance in grapevine downy mildew populations. Appl Environ Microbiol. 2007;73(16): 5162–5172.

7. Ballabio C, Panagos P, Lugato E, Huang J-H, Orgiazzi A, Jones A, et al. Copper distribution in European topsoils: An assessment based on LUCAS soil survey. Sci Total Environ. 2018;636: 282–298.

8. Flemming CA, Trevors JT, Pollution S. Copper toxicity and chemistry in the environment: a review. Water Air Soil Pollut. 1989;44(1-2):143–158.

9. Salmon JM, Friedl MA, Frolking S, Wisser D, Douglas EM. Global rain-fed, irrigated, and paddy croplands: A new high resolution map derived from remote sensing, crop inventories and climate data. Int J Appl Earth Obs Geoinf. 2015;38:321–334.

10. Viret O, Spring JL, Gindro K. Stilbenes: biomarkers of grapevine resistance to fungal diseases. OENO One. 2018;52(3): 235–241.

11. Yobrégat O. Introduction to resistant vine types: a brief history and overview of the situation. OENO One. 2018;52: 241–246.

12. De la Fuente Lloreda M. Use of hybrids in viticulture. A challenge for the OIV. OENO One. 2018;52(3): 231–234.

13. Peressotti E, Wiedemann-Merdinoglu S, Delmotte F, Bellin D, Di Gaspero G, Testolin R, et al. Breakdown of resistance to grapevine downy mildew upon limited deployment of a resistant variety. BMC Plant Biol. 2010;10(1): 147.

14. Boller T, Felix G. A renaissance of elicitors: perception of microbe-associated molecular patterns and danger signals by pattern-recognition receptors. Annu Rev Plant Biol. 2009;60: 379–406.

15. Yi SY, Shirasu K, Moon JS, Lee S-G, Kwon S-Y. The activated SA and JA signaling pathways have an influence on flg22-triggered oxidative burst and callose deposition. PloS one. 2014;9(2): e88951-e.

16. Gomès E, Coutos-Thévenot P. Molecular Aspects of grapevine-pathogenic fungi interactions. In: Roubelakis-Angelakis KA, editor. Grapevine Molecular Physiology & Biotechnology. Dordrecht: Springer Netherlands; 2009. p. 407–428.

17. Bektas Y, Eulgem T. Synthetic plant defense elicitors. Front Plant Sci. 2015;5: 804.

18. Hamiduzzaman MM, Jakab G, Barnavon L, Neuhaus JM, Mauch-Mani B. beta-aminobutyric acid-induced resistance against downy mildew in grapevine acts through the potentiation of callose formation and jasmonic acid signaling. Mol Plant Microbe Iinteract. 2005;18(8): 819–689.

19. Aziz A, Trotel-Aziz P, Dhuicq L, Jeandet P, Couderchet M, Vernet G. Chitosan oligomers and copper sulfate induce grapevine defense reactions and resistance to gray mold and downy mildew. Phytopathol. 2006;96(11):c1188–1194.

20. Trouvelot S, Varnier AL, Allegre M, Mercier L, Baillieul F, Arnould C, et al. A beta-1,3 glucan sulfate induces resistance in grapevine against *Plasmopara viticola* through priming of defense responses, including HR-like cell death. Mol Plant Microbe Interact. 2008;21(2): 232–243.

21. Allegre M, Heloir MC, Trouvelot S, Daire X, Pugin A, Wendehenne D, et al. Are grapevine stomata involved in the elicitor-induced protection against downy mildew? Mol Plant Microbe Inter. 2009;22(8): 977–986.

22. Delaunois B, Farace G, Jeandet P, Clement C, Baillieul F, Dorey S, et al. Elicitors as alternative strategy to pesticides in grapevine? Current knowledge on their mode of action from controlled conditions to vineyard. Environ Sci Poll Res Int. 2013;21(7): 4837–4846.

23. Godard S, Slacanin I, Viret O, Gindro K. Induction of defence mechanisms in grapevine leaves by emodin- and anthraquinone-rich plant extracts and their conferred resistance to downy mildew. Plant Physiol Biochem. 2009;47(9): 827–837.

24. Dagostin S, Formolo T, Giovannini O, Pertot I. *Salvia officinalis* extract can protect grapevine against *Plasmopara viticola*. Plant Dis. n°5. 2010;94(36): 575–580.

25. Soylu EM, Kurt Ş, Soylu S. In vitro and in vivo antifungal activities of the essential oils of various plants against tomato grey mould disease agent Botrytis cinerea. Int J Food Microbiol. 2010;143(3): 183–189.

26. Pina-Vaz C, Goncalves Rodrigues A, Pinto E, Costa-de-Oliveira S, Tavares C, Salgueiro L, et al. Antifungal activity of *Thymus* oils and their major compounds. J Eur Acad Dermatol Venereol. 2004;18(1): 73–78.

27. Soylu S, Yigitbas H, Soylu EM, Kurt S. Antifungal effects of essential oils from oregano and fennel on *Sclerotinia sclerotiorum*. J Appl Microbiol. 2007;103(4): 1021–1030.

28. Soylu EM, Kurt S, Soylu S. In vitro and in vivo antifungal activities of the essential oils of various plants against tomato grey mould disease agent *Botrytis cinerea*. Int J Food Microbiol. 2010;143(3): 183–189.

29. Dagostin S, Formolo T, Giovannini O, Pertot I, Schmitt A. *Salvia officinalis* extract can protect grapevine against *Plasmopara viticola*. Plant Dis. 2010;94(5): 575–580.

30. Turek C, Stintzing FC. Stability of essential oils: A review. Compr Rev Food Sci Food Saf. 2013;12(1): 40–53.

31. Janatova A, Bernardos A, Smid J, Frankova A, Lhotka M, Kourimská L, et al. Long-term antifungal activity of volatile essential oil components released from mesoporous silica materials. Ind Crops Prod. 2015;67: 2216–20.

32. Oliveira Fialho R, Stradioto Papa MdF, Rodrigo Panosso A, Rodrigues Cassiolato A. Fungitoxicity of essential oils on *Plasmopora viticola*, causal agent of grapevine downy mildew. Rev Bras Frutic. 2017; 39(4): e-015.

33. Burketova L, Trda L, Ott PG, Valentova O. Bio-based resistance inducers for sustainable plant protection against pathogens. Biotechnol Adv. 2015;33(6 Pt 2): 994–1004.

34. Teixeira B, Marques A, Ramos C, Serrano C, Matos O, Neng NR, et al. Chemical composition and bioactivity of different oregano (Origanum vulgare) extracts and essential oil. J Sci Food Agric. 2013;93(11): 2707–2714.

35. Bozin B, Mimica-Dukic N, Simin N, Anackov G. Characterization of the volatile composition of essential oils of some Lamiaceae spices and the antimicrobial and antioxidant activities of the entire oils. J Agric Food Chem. 2006;54(5): 1822–188.

36. Bisht D, Chanotiya CS, Rana M, Semwal M. Variability in essential oil and bioactive chiral monoterpenoid compositions of Indian oregano (*Origanum vulgare* L.) populations from northwestern Himalaya and their chemotaxonomy. Ind Crops Prod. 2009;30(3): 42422–6.

37. Boruga O, Jianu C, Misca C, Golet I, Gruia AT, Horhat FG. *Thymus vulgaris* essential oil: chemical composition and antimicrobial activity. J Med Life. 2014;7(Spec 3): 56–60.

38. Sakkas H, Papadopoulou C. Antimicrobial Activity of basil, oregano, and thyme essential oils. J iMcrobiol Biotechnol. 2017;27(3): 429–438.

39. Mohamed A., Hamza A., A D. Recent approaches for controlling downy mildew of cucumber under greenhouse conditions. Plant Protect Sci. 2016;52: 1–9.

40. Nowak A, Konstantinidou-Doltsinis S, Seddon B, Schmitt A. Alternative agents for control of downy mildew (*Pseudoperonospora cubensis*) of cucumbers. Mitt Julius Kühn-Inst. 2008(419): 64–67.

41. La Torre A, Mandalà C, Pezza L, Caradonia F, Battaglia V. Evaluation of essential plant oils for the control of Plasmopara viticola. J Essent Oil Res. 2014;26(4):c282–291.

42. Nazzaro F, Fratianni F, Coppola R, Feo VD. Essential oils and antifungal activity. Pharmaceuticals. 2017;10(4): 86.

43. Nobrega RO, Teixeira AP, Oliveira WA, Lima EO, Lima IO. Investigation of the antifungal activity of carvacrol against strains of *Cryptococcus neoformans*. Pharm Biol. 2016;54(11): 2591–2596.

44. Chavan PS, Tupe SG. Antifungal activity and mechanism of action of carvacrol and thymol against vineyard and wine spoilage yeasts. Food Contr. 2014;46: 115–120.

45. Yin L, An Y, Qu J, Li X, Zhang Y, Dry I, et al. Genome sequence of *Plasmopara viticola* and insight into the pathogenic mechanism. Sci Rep. 2017;7: 46553.

46. Broekaert WF, Delauré SL, Bolle MFCD, Cammue BPA. The role of ethylene in host-pathogen interactions. Annu Rev Phytopathol. 2006;44(1): 393–416.

47. Pieterse CMJ, Does DVd, Zamioudis C, Leon-Reyes A, Wees SCMV. Hormonal modulation of plant immunity. Annu Rev Cell Dev Biol. 2012;28(1): 489–521.

48. Santino A, Taurino M, De Domenico S, Bonsegna S, Poltronieri P, Pastor V, et al. Jasmonate signaling in plant development and defense response to multiple (a)biotic stresses. Plant Cell Rep. 2013;32(7): 1085–1098.

49. Larrieu A, Vernoux T. Q&A: How does jasmonate signaling enable plants to adapt and survive? BMC biology. 2016; 14:79.

50. Boubakri H, Wahab MA, Chong J, Bertsch C, Mliki A, Soustre-Gacougnolle I. Thiamine induced resistance to *Plasmopara viticola* in grapevine and elicited host-defense responses, including HR like-cell death. Plant Physiol Biochem. 2012;57:120–133.

51. Pajerowska-Mukhtar KM, Emerine DK, Mukhtar MS. Tell me more: roles of NPRs in plant immunity. Trends Plant Sci. 2013;18(7): 402–411.

52. Spoel SH, Koornneef A, Claessens SMC, Korzelius JP, Van Pelt JA, Mueller MJ, et al. NPR1 Modulates cross-talk between salicylate- and jasmonate-dependent defense pathways through a novel function in the cytosol. 2003;15(3):760–70.

53. Backer R, Naidoo S, van den Berg N. The nonexpressor of pathogenesis-related genes 1 (NPR1) and related family: mechanistic insights in plant disease resistance. Front Plant Sci. 2019;10: 102.

54. Chini A, Fonseca S, Fernandez G, Adie B, Chico JM, Lorenzo O, et al. The JAZ family of repressors is the missing link in jasmonate signalling. Nature. 2007;448(7154): 666–671.

55. Kazan K, Manners JM. JAZ repressors and the orchestration of phytohormone crosstalk. Trends Plant Sci. 2012;17(1): 22–31.

56. Proietti S, Caarls L, Coolen S, Van Pelt JA, Van Wees SCM, Pieterse CMJ. Genome-wide association study reveals novel players in defense hormone crosstalk in *Arabidopsis*. Plant Cell Environment. 2018;41(10): 2342–2356.

57. Caarls L, Pieterse CMJ, Van Wees SCM. How salicylic acid takes transcriptional control over jasmonic acid signaling. Front Plant Sci. 2015;6: 170.

58. Zheng Z, Qamar SA, Chen Z, Mengiste T. Arabidopsis WRKY33 transcription factor is required for resistance to necrotrophic fungal pathogens. Plant J. 2006;48(4): 592–605.

59. Li J, Brader G, Kariola T, Palva ET. WRKY70 modulates the selection of signaling pathways in plant defense. Plant J. 2006;46(3): 477–491.

60. Wang D, Amornsiripanitch N, Dong X. A genomic approach to identify regulatory nodes in the transcriptional network of systemic acquired resistance in plants. PLoS Pathog. 2006;2(11): e123.

61. Pandey SP, Somssich IE. The Role of WRKY Transcription Factors in Plant Immunity. Plant Physiol. 2009;150(4): 1648–1655.

62. Paul PK, Sharma PD. *Azadirachta indica* leaf extract induces resistance in barley against leaf stripe disease. Physiol Mol Plant Pathol. 2002;61(1): 3–13.

63. Gao Q-M, Zhu S, Kachroo P, Kachroo A. Signal regulators of systemic acquired resistance. Front Plant Sci. 2015;6: 228.

64. Vergnes S, Ladouce N, Fournier S, Ferhout H, Attia F, Dumas B. Foliar treatments with Gaultheria procumbens essential oil induce defense responses and resistance against a fungal pathogen in Arabidopsis. 2014; 5: 477.

65. von Rad U, Mueller MJ, Durner J. Evaluation of natural and synthetic stimulants of plant immunity by microarray technology. New Phytol. 2005;165(1): 191–202.

66. Gullner G, Komives T, Kiraly L, Schroder P. Glutathione S-Transferase enzymes in plant-pathogen interactions. Front Plant Sci. 2018;9: 1836.

67. Suzuki N, Sejima H, Tam R, Schlauch K, Mittler R. Identification of the MBF1 heat-response regulon of *Arabidopsis thaliana*. Plant J. 2011;66: 844–851.

68. Suzuki N, Koussevitzky S, Mittler R, Miller G. ROS and redox signalling in the response of plants to abiotic stress. Plant Cell Environ. 2012;35(2): 259–270.

69. Rienth M, Torregrosa L, Sarah G, Ardisson M, Brillouet J-M, Romieu C. Temperature desynchronizes sugar and organic acid metabolism in ripening grapevine fruits and remodels their transcriptome. BMC Plant Biol. 2016;16(1): 164.

70. Rienth M, Torregrosa L, Kelly MT, Luchaire N, Pellegrino A, Grimplet Jrm, et al. Is transcriptomic regulation of berry development more important at night than during the day? PLoS One. 2014;9(2): e88844.

71. Rienth M, Torregrosa L, Luchaire N, Chatbanyong R, Lecourieux D, Kelly M, et al. Day and night heat stress trigger different transcriptomic responses in green and ripening grapevine (*Vitis vinifera*) fruit. BMC Plant Biol. 2014;14(1): 108.

72. Mizoi J, Shinozaki K, Yamaguchi-Shinozaki K. AP2/ERF family transcription factors in plant abiotic stress responses. Biochim Biophys Acta. - Gene Regulatory Mechanisms. 2012;1819(2):86–96.

73. Bahieldin A, Atef A, Edris S, Gadalla NO, Ali HM, Hassan SM, et al. Ethylene responsive transcription factor ERF109 retards PCD and improves salt tolerance in plant. BMC Plant Biol. 2016;16(1): 216.

74. Kumar M, Brar A, Yadav M, Chawade A, Vivekanand V, Pareek N. Chitinases-potential candidates for enhanced plant resistance towards fungal pathogens. Agriculture 2018, 8(7), 88.

75. Hakim, Ullah A, Hussain A, Shaban M, Khan AH, Alariqi M, et al. Osmotin: A plant defense tool against biotic and abiotic stresses. Plant Physiol Biochem. 2018;123: 149–159.

76. Gauthier A, Trouvelot S, Kelloniemi J, Frettinger P, Wendehenne D, Daire X, et al. The sulfated laminarin triggers a stress transcriptome before priming the SA- and ROS-dependent defenses during grapevine’s induced resistance against *Plasmopara viticola*. PLOS ONE. 2014;9(2): e88145.

77. Rasmussen M, Roux M, Petersen M, Mundy J. MAP Kinase Cascades in *Arabidopsis* Innate Immunity. Front Plant Sci. 2012;3: 169.

78. Tsuda K, Mine A, Bethke G, Igarashi D, Botanga CJ, Tsuda Y, et al. Dual regulation of gene expression mediated by extended MAPK activation and salicylic acid contributes to robust innate immunity in *Arabidopsis thaliana*. PLoS Genet. 2013;9(12): e1004015.

79. Su J, Yang L, Zhu Q, Wu H, He Y, Liu Y, et al. Active photosynthetic inhibition mediated by MPK3/MPK6 is critical to effector-triggered immunity. PLOS Biol. 2018;16(5): e2004122.

80. Shoresh M, Gal-On A, Leibman D, Chet I. Characterization of a mitogen-activated protein kinase gene from cucumber required for *Trichoderma*-conferred plant resistance. Plant Physiol. 2006;142(3): 1169–1179.

81. Genot B, Lang J, Berriri S, Garmier M, Gilard F, Pateyron S, et al. Constitutively active *Arabidopsis* MAP Kinase 3 triggers defense responses involving salicylic acid and SUMM2 resistance protein. Plant Physiol. 2017;174(2): 1238–1249.

82. Kwaaitaal M, Huisman R, Maintz J, Reinstädler A, Panstruga R. Ionotropic glutamate receptor (iGluR)-like channels mediate MAMP-induced calcium influx in *Arabidopsis thaliana*. Biochem J. 2011;440(3): 355–365.

83. CyanoBase. http://genome.microbedb.jp/cyanobase/

84. Zhu X, Caplan J, Mamillapalli P, Czymmek K, Dinesh-Kumar SP. Function of endoplasmic reticulum calcium ATPase in innate immunity-mediated programmed cell death. EMBO J. 2010;29(5): 1007–1018.

85. Lam E, Kato N, Lawton M. Programmed cell death, mitochondria and the plant hypersensitive response. Nature. 2001;411(6839): 848–853.

86. Stael S, Kmiecik P, Willems P, Van Der Kelen K, Coll NS, Teige M, et al. Plant innate immunity--sunny side up? Trends Plant Sci. 2015;20(1): 3–11.

87. Anderson JP, Badruzsaufari E, Schenk PM, Manners JM, Desmond OJ, Ehlert C, et al. Antagonistic interaction between abscisic acid and jasmonate-ethylene signaling pathways modulates defense gene expression and disease resistance in *Arabidopsis*. Plant Cell. 2004;16(12): 3460–3479.

88. Vanlerberghe GC. Alternative oxidase: a mitochondrial respiratory pathway to maintain metabolic and signaling homeostasis during abiotic and biotic stress in plants. Int J Mol Sci. 2013;14(4): 6805–6847.

89. Piasecka A, Jedrzejczak-Rey N, Bednarek P. Secondary metabolites in plant innate immunity: conserved function of divergent chemicals. New Phytol. 2015;206(3): 948–964.

90. Mierziak J, Kostyn K, Kulma A. Flavonoids as important molecules of plant interactions with the environment. Molecules. 2014;19(10): 16240–16265.

91. Fofana B, McNally DJ, Labbé C, Boulanger R, Benhamou N, Séguin A, et al. Milsana-induced resistance in powdery mildew-infected cucumber plants correlates with the induction of chalcone synthase and chalcone isomerase. Physiol Mol Plant Pathol. 2002;61(2):1 21–32.

92. Richard T, Abdelli-Belhad A, Vitrac X, Waffo-Téguo P, Mérillon J-M. *Vitis vinifer*a canes, a source of stilbenoids against downy mildew. OENO One. 2016;50(3).

93. Höll J, Vannozzi A, Czemmel S, D’Onofrio C, Walker AR, Rausch T, et al. The R2R3-MYB Transcription Factors MYB14 and MYB15 Regulate Stilbene Biosynthesis in Vitis vinifera. 2013;25(10): 4135–4149.

94. Belhadj A, Telef N, Saigne C, Cluzet S, Barrieu F, Hamdi S, et al. Effect of methyl jasmonate in combination with carbohydrates on gene expression of PR proteins, stilbene and anthocyanin accumulation in grapevine cell cultures. Plant Physiol Biochem. 2008;46: 493–499.

95. D’Onofrio C, Cox A, Davies C, Boss P. Induction of secondary metabolism in grape cell cultures by jasmonates. Funct Plant Biol. 2009;36: 323–338.

96. Garde-Cerdan T, Gutiérrez-Gamboa, G. P-Á, Pérez-Álvarez E. P., Rubio-Bretón P. Foliar application of methyl jasmonate to Graciano and Tempranillo vines: effects on grape amino acid content during two consecutive vintages. OENO One,. 2019; 53(1).

97. Kordali S, Cakir A, Ozer H, Cakmakci R, Kesdek M, Mete E. Antifungal, phytotoxic and insecticidal properties of essential oil isolated from Turkish *Origanum acutidens* and its three components, carvacrol, thymol and p-cymene. Biores Technol. 2008;99(18): 8788–8795.

98. de Almeida LFR, Frei F, Mancini E, De Martino L, De Feo V. Phytotoxic activities of Mediterranean essential oils. Molecules. 2010;15(6): 4309–4323.

99. Amri IS, Hamrouni L, Hananac M, Jamoussi B. Reviews on phytotoxic effects of essential oils and their individual components: news approach for weeds management. Int J Appl BiolPharm Technol. 2012: 96–114.

100. Duchêne E, Dumas V, Jaegli N, Merdinoglu D. Genetic variability of descriptors for grapevine berry acidity in Riesling, Gewürztraminer and their progeny. Aust J Grape Wine Res. 2014;20(1): 91–99.

101. Merdinoglu D, Schneider C, Prado E, Wiedemann-Merdinoglu S, Mestre P. Breeding for durable resistance to downy and powdery mildew in grapevine. OENO One. 2018;52(3): 103–209.

102. Torregrosa L, Bigard A, Doligez A, Lecourieux D, Rienth M, Luchaire N, et al. Developmental, molecular and genetic studies on grapevine response to temperature open breeding strategies for adaptation to warming. OENO One. 2017;51(2): 155–165.

103. Rienth M, Torregrosa L, Ardisson M, De Marchi R, Romieu C. Versatile and efficient RNA extraction protocol for grapevine berry tissue, suited for next generation RNA sequencing. Aust J Grape Wine Res. 2014;20(2): 247–254.

104. Leinonen R, Sugawara H, Shumway M. The sequence read archive. Nucleic Acids Res. 2011;39: D19–21.

105. Kim D, Langmead B, Salzberg SL. HISAT: a fast spliced aligner with low memory requirements. Nat Methods. 2015;12(4): 357–360.

106. Anders S, Pyl PT, Huber W. HTSeq--a Python framework to work with high-throughput sequencing data. Bioinformatics. 2015;31(2): 166–159.

107. Love MI, Huber W, Anders S. Moderated estimation of fold change and dispersion for RNA-seq data with DESeq2. Genome Biol. 2014;15(12): 550.

108. Al-Shahrour F, Diaz-Uriarte R, Dopazo J. FatiGO: a web tool for finding significant associations of gene ontology terms with groups of genes. Bioinformatics. 2004;20(4): 578–580.

109. Grimplet J, Van Hemert J, Carbonell-Bejerano P, Diaz-Riquelme J, Dickerson J, Fennell A, et al. Comparative analysis of grapevine whole-genome gene predictions, functional annotation, categorization and integration of the predicted gene sequences. BMC Res Notes. 2012;5(1): 213.

